# TailEnrich enables discovery of dysregulated genes through disease sample enrichment testing

**DOI:** 10.64898/2026.11.04.764457

**Authors:** Hongdong Li, Yuhe Liu, Cuixiang Lin, Shixian Wang, Hui Pan, Shaokai Wang

## Abstract

Identifying differentially expressed genes is fundamental to elucidating the molecular mecha nisms underlying complex diseases. However, in complex disorders such as Alzheimer’s disease (AD), the genes exhibiting dysregulated expression may vary among patient subsets, reflecting the heterogeneity of the underlying pathological processes. Consequently, pathogenic genes are profoundly dysregulated in only a subset of patients. Conventional differential expression tools, such as DESeq2 and edgeR, are primarily designed to detect population-level expression shifts and may have limited power to identify subgroup-specific dysregulation. To overcome this limitation, we developed TailEnrich, a novel com putational method that discovers dysregulated genes within patient subsets. TailEnrich independently tests the extreme left (downregulated) or right (upregulated) tails of a gene’s expression distribution for disease sample enrichment. We quantify this enrichment using a new metric, i.e. the PE (Percentage of Enriched disease samples) score. On simulated datasets, TailEnrich demonstrates markedly improved accuracy over DESeq2 and edgeR in identifying dysregulated genes specific to patient subsets. Applied to AD gene expression data, TailEnrich uncovers a distinct, biologically relevant set of genes within key pathological pathways that are missed by mean shift-based approaches. TailEnrich provides a novel method for identifying dysregulated genes, and readily extends to other omics data including genomic variants and proteomics to uncover pathogenic genes in complex diseases.

**Availability and implementation:** TailEnrich is implemented in R. The source code and documenta tion are freely available at https://github.com/Sanke926/Tailenrich.

## 1 Introduction

Identifying differentially expressed genes is fundamental to elucidating the molecular mechanisms underlying complex diseases. However, complex diseases, such as Alzheimer’s disease (AD), exhibit profound molecular heterogeneity and often encompass different dysregulated genes driven by distinct pathologies [1]. With a single clinical diagnosis, different patients may be governed by divergent mechanisms, such as neuroinflamma tion versus lipid dysregulation in AD [2]. Due to the large difference in pathological drivers, gene expression changes are rarely uniform across the entire patient population. Consequently, pathogenic genes may be profoundly dysregulated in only a specific subset of patients. If a gene is dysregulated in only a subset of patients, the corresponding expression abnormality may be concentrated in that subgroup and diluted at the population level. Capturing subset-specific dysregulation is therefore important for describing the molecular heterogeneity of complex diseases [3].

Conventional differential expression tools, most notably DESeq2 and edgeR (v4), have become the stan dard for transcriptomic analysis [4, 5], but they fundamentally rely on detecting population-level mean shifts of gene expression. These powerful algorithms model RNA sequencing count data using a negative binomial distribution, estimating dispersion to account for biological variance, and ultimately testing for significant differences in the average gene expression between all patients and all controls [4, 5]. However, these mean-centric approaches are challenged when analyzing gene expression data of complex diseases. If a pathogenic gene is heavily dysregulated in only a small subset of patients, the baseline expression of that gene in the unaffected majority of the disease cohort mathematically dilutes the signal. When the algorithm calculates the population mean, the extreme low or high expression values from the subset of patients are averaged out, rendering the overall disease group statistically indistinguishable from the control group. As a result, these methods can miss genes that are dysregulated only in a subset of patients [3].

To overcome the limitation of mean-shift dilution, we developed TailEnrich, a novel computational method specifically designed to discover dysregulated genes hidden within patient subsets. Rather than focusing on mean changes of gene expression, TailEnrich adopts a tail-centric framework. The core innovation lies in independently testing the extreme left (downregulated) or right (upregulated) tail of a gene’s expression distribution for disease sample enrichment. By scanning the expression profile of each gene, TailEnrich identifies genes where a disproportionate number of disease samples occupy the extreme expression deciles. To quantify this enrichment, we introduce a new, interpretable metric: the PE (Percentage of Enriched disease samples) score. The significance of the PE score is evaluated through a permutation test. By isolating signals at the extreme end of the gene expression distribution, this approach is capable of detecting genes that are dysregulated only in a subset of patients. We test the validity of TailEnrich across both simulated and real-world disease data. Using simulated datasets modeled to reflect disease heterogeneity, TailEnrich demonstrates markedly improved accuracy and sensitivity over DESeq2 and edgeR (v4) in capturing patient subset-specific dysregulation. We then applied TailEnrich to RNA-seq gene expression data from AD cohorts. TailEnrich identified a distinct set of AD-associated genes enriched in relevant pathways that were not detected by mean-based approaches. Ultimately, TailEnrich provides a novel complement to conventional mean-based approaches, expanding the landscape of pathogenic gene discovery in complex, heterogeneous diseases.

## 2 Methods

TailEnrich is a computational method designed to identify genes whose expression is enriched for disease samples at the low- or high-expression tails across samples. Unlike conventional differential analysis methods that primarily test mean differences between disease and control samples, TailEnrich evaluates whether disease samples are preferentially enriched in either the low-expression or high-expression tail of each gene. This design is intended to identify disease subset-specific dysregulated genes that are likely to be missed by conventional approaches.

### 2.1 Input

The input of TailEnrich consists of a gene expression matrix and the corresponding sample labels (Fig. 1a). For a dataset containing *m* genes and *n* samples, let **X** denote an *m × n* gene expression matrix, in which each row represents a gene and each column represents a sample. The expression profile of a gene across all samples is therefore represented by one row of **X**. Let **y** denote an *n*-dimensional label vector corresponding to the samples in **X**, where *y* = +1 indicates a disease sample and *y* = *−*1 indicates a control sample.

**Fig 1.**
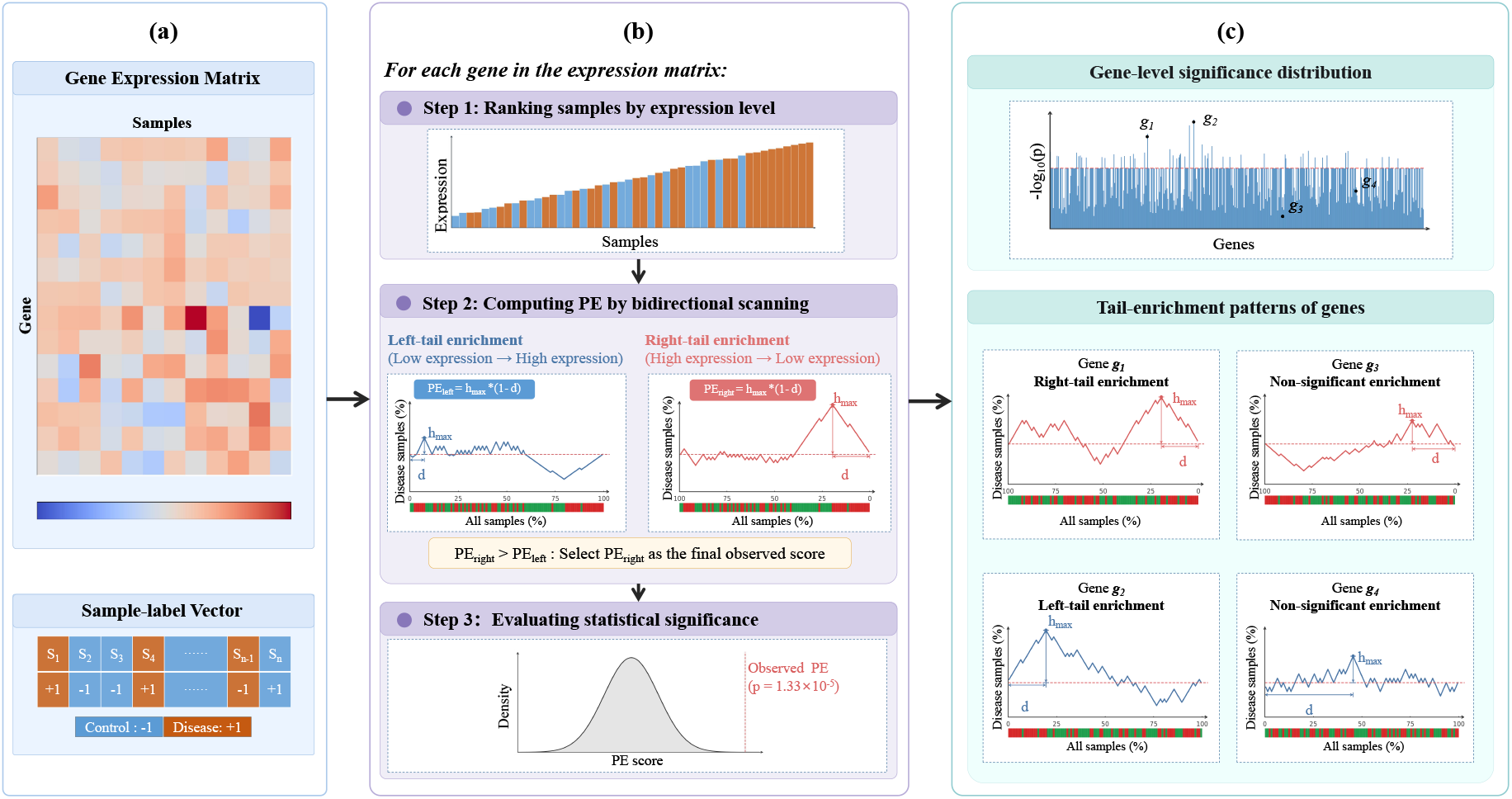
Workflow of TailEnrich. **(a)** TailEnrich takes a gene expression matrix and a sample-label vector as input. Control and disease samples are encoded as *−* 1 and +1, respectively. **(b)** For each gene, samples are first ranked by expression level. TailEnrich then computes PE for two directions, corresponding to left-tail and right-tail enrichment. The color strip below each curve indicates the sample group at each ranked position; green and red denote control and disease samples, respectively. The larger of the two PE values is retained as the final PE. Statistical significance is evaluated by permutation testing, and the Benjamini–Hochberg procedure is applied to calculate FDR. **(c)** An illustrative example of the distribution of gene *p* values by TailEnrich. Four example genes along with their enrichment patterns are highlighted: *g*_1_ shows right-tail enrichment, *g*_2_ shows left-tail enrichment, and *g*_3_ and *g*_4_ show no significant tail enrichment.

### 2.2 The TailEnrich algorithm

The workflow of TailEnrich is shown in Fig. 1. It consists of three main steps: sample ranking, calculation of the weighted Percentage of Enriched (PE) disease samples, and significance test. For each gene, TailEnrich ranks all samples by expression values in ascending order to calculate the left-tail and right-tail PE scores. The larger PE score is retained as the final score, with its statistical significance subsequently evaluated via permutation testing.

#### Ranking samples by gene expression level

For each gene, TailEnrich sorts all samples by expression level in ascending order. The sorted samples are next used to compute the enrichment of the gene in disease samples.

#### Computing PE score for each gene

Before enrichment analysis, sample labels are first normalized. Let *yi* denote the sample label, with +1 and *−* 1 representing disease and control samples, respectively, which is normalzied using the formula:

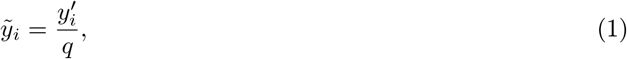

where 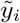 denotes the normalized label, 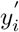 denotes the centered label, and *q* denotes the sum of the centered labels of all disease samples. In detail, the centered label 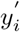 is calculated by substracting the mean of all sample labels from *yi*. In this normalization, centering is used to account for unbalanced sample sizes between disease and control groups, and dividing by *q* is used to make the maximum percentage of enriched disease samples (i.e. *h*max below) equal to 100% for the convenience of understanding.

TailEnrich calculates a PE score independently for left-tail and right-tail enrichment. PE is defined as the weighted Percentage of Enriched disease samples:

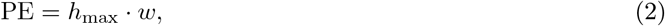

where *h*max denotes the maximum percentage of enriched disease samples, and *w* is a weight of which the value is related to the location of this maximum. The process for calculating the PE score of a gene is described in three steps below.

1. Left-tail enrichment score (PEleft) calculation. Let 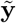 denote the normalized sample-label vector of the sorted samples. The percentage of enriched disease samples across the ranked positions is represented by a vector **h** = (*h*1, …, *hn*)^⊤^ and is calculated as

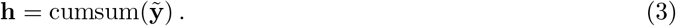

The cumsum denotes the operation of calculating the cumulative sum of the normalized sample labels. As an example, the *i*th element of **h** is calculated as

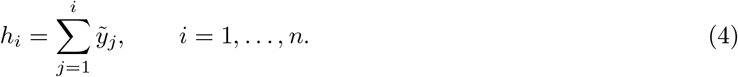

The maximum of the percentage of enriched disease samples, denoted by *h*max, is calculated as

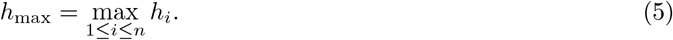

Let *d* denote the percentage of the samples whose gene expression level is equal to or lower than the expression of the *k*th sample, where *k* is the sample index at which the percentage of enriched disease samples achieves the maximum. Then, the corresponding weight is calculated as

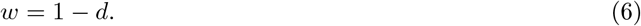

The left-tail PE is then calculated as

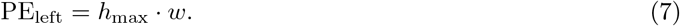

1. Right-tail enrichment score (PEright) calculation. For right-tail enrichment, samples are analyzed from high-to-low expression end. The percentage of enriched disease samples h and its maximum *h*max can be calculated as in Eqs. 3–5 but in the high-to-low expression direction (i.e. from right to left). Let *d* denote the percentage of samples whose gene expression level is equal to or higher than the expression of the *k*th sample, where *k* is the sample index at which the percentage of enriched disease samples achieves the maximum. Then, the corresponding weight is calculated as *w* = 1 *− d*.

The right-tail PE is then calculated as

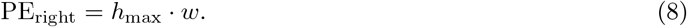

1. Determination of the PE score of the gene. Because the enriched tail of a given gene is unknown in advance, TailEnrich calculates PE independently in both directions as described above and retains the larger of PEleft and PEright as the final PE for each gene:

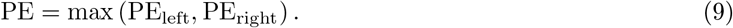

Specifically, in the extreme case where the expression distributions of the disease and control samples are completely separated and the two groups are equal in size, we will have *h*max = 100%, *w* = 0.5, and PE = 0.5 (as illustrated in Supplementary Figure S1).

#### Evaluating statistical significance by permutation test

TailEnrich evaluates the statistical significance of the final observed PE through a permutation test (Fig. 1b, Step 3). For each permutation, the sample labels are randomly permuted while keeping the gene expression data unchanged, and the PE score of a given gene is calculated based on the permuted data. The PE values resulting from *N*perm permutations are used as the null distribution. For each gene, the *p* value of its PE score is calculated as

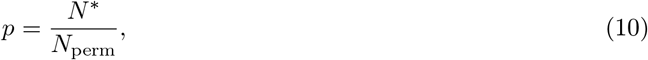

where *N* ^*\**^ denotes the number of permuted PE scores that are equal to or greater than the observed PE of the gene. Finally, the resulting *p* values of all genes are adjusted using the Benjamini–Hochberg procedure [6] and reported as false discovery rate (FDR). The resulting gene-level significance distribution and representative tail-enriched genes with left-tail or right-tail enrichment are illustrated in Fig. 1c.

## 3 Results and discussion

We evaluated TailEnrich using both simulated data and real-world Alzheimer’s disease (AD) RNA-seq data. TailEnrich was compared with the two most widely used differential analysis methods, DESeq2 [4] and edgeR (v4) [5].

### 3.1 TailEnrich outperforms DESeq2 and edgeR in simulation studies

We compared TailEnrich with DESeq2 and edgeR on simulated datasets under two types of tail enrichment: (I) left-tail enrichment, in which a subset of disease samples was enriched at the left (low-expression) tail of the gene expression distribution; and (II) right-tail enrichment, in which a subset of disease samples was enriched at the right (high-expression) tail of the gene expression distribution.

The simulated datasets were generated in two stages. First, a baseline expression matrix without differen tial expression was generated using the generateSyntheticData function in the compcodeR package (version 1.38.0) [7], a widely used approach to simulate RNA-seq data. Each simulated dataset contained 5,000 genes and 100 samples, including 50 disease samples and 50 control samples. Second, in tail enrichment simulation, 10% of the genes (i.e. 500 genes) were set to show enrichment at the tail of the gene expression distribution. Two parameters, *p* and *r*, were used to control the simulated data. The parameter *p* specified the proportion of disease samples enriched at the tail of the gene expression distribution. The parameter *r* denoted the degree of dysregulation. A larger *r* value indicated that the gene expression of disease samples enriched in the tail had a larger difference compared to control samples. In this study, *p* was set to 20%, 30%, or 40%, and *r* was set to 0.1, 0.3, or 0.5. In detail, to simulate a dysregulated gene, its expression in a proportion (*p*) of disease samples was modified to be lower (left-tail enrichment) or higher (right-tail enrichment) than that in all remaining samples. The detailed data-simulation procedures and parameter settings are provided in Supplementary Note S1 and Supplementary Table S1.

TailEnrich better distinguished genes dysregulated in a subset of disease samples than DESeq2 and edgeR. We applied the three methods to the simulated datasets described above and ranked the genes according to the *−*log10(*p*) value. For each parameter setting, their performance in distinguishing the 500 tail-enriched genes from the remaining 4,500 background genes was evaluated using AUROC and AUPRC over 20 independent replicates. Considering left-tail and right-tail enrichment and all values of *p* and *r*, the simulation study included 18 parameter combinations. For AUROC, TailEnrich achieved the highest value in 10 of the 18 parameter combinations. For AUPRC, it achieved the highest value in 16 of the 18 parameter combinations. The mean AUROC values of TailEnrich ranged from 0.953 to 1.000. Its mean AUPRC values ranged from 0.877 to 1.000 (Table 1). The detailed results for TailEnrich, edgeR, and DESeq2 are provided in Supplementary Tables S2–S4, respectively. Performance distributions for the three methods across all parameter settings in simulations are shown in Supplementary Figures S2–S3. For example, when *p* = 30%, TailEnrich maintained relatively high AUROC and AUPRC values under both left-tail and right-tail enrichment across different values of *r* (Fig. 2a,b). Taking the setting of *p* = 30% and *r* = 0.3 as an example, the AUROC values of TailEnrich, DESeq2, and edgeR under left-tail enrichment were 0.994 *±* 0.008, 0.993 *±* 0.011, and 0.997 *±* 0.006, respectively (Fig. 2a,c). Their AUPRC values were 0.984 *±* 0.021, 0.937*±* 0.076, and 0.975 *±* 0.046, respectively (Fig. 2a,c). Under right-tail enrichment, the AUROC values of TailEnrich, DESeq2, and edgeR were 0.996 *±* 0.006, 0.980 *±* 0.016, and 0.991 *±* 0.009, respectively (Fig. 2b,d). Their AUPRC values were 0.992 *±* 0.011, 0.855 *±* 0.094, and 0.940 *±* 0.060, respectively (Fig. 2b,d). We also conducted additional simulations with 10,000 genes, and performance distributions of the three methods are shown in Supplementary Figures S4–S5. Following the method described in Wu et al. [8], we further compared the recall value of the three methods at different low FDR levels under the same 18 parameter combinations as above. The complete FDR–Recall curves for all parameter combinations are shown in Supplementary Figures S6–S9. Taking *FDR* = 0.05 as an example, TailEnrich achieved the highest mean recall in 16 of the 18 parameter combinations. Specifically, when *p* = 30% and *r* = 0.3, TailEnrich achieved higher mean recall than DESeq2 and edgeR at all FDR levels under both left-tail and right-tail enrichment (Fig. 2c,d).

**Table 1.**
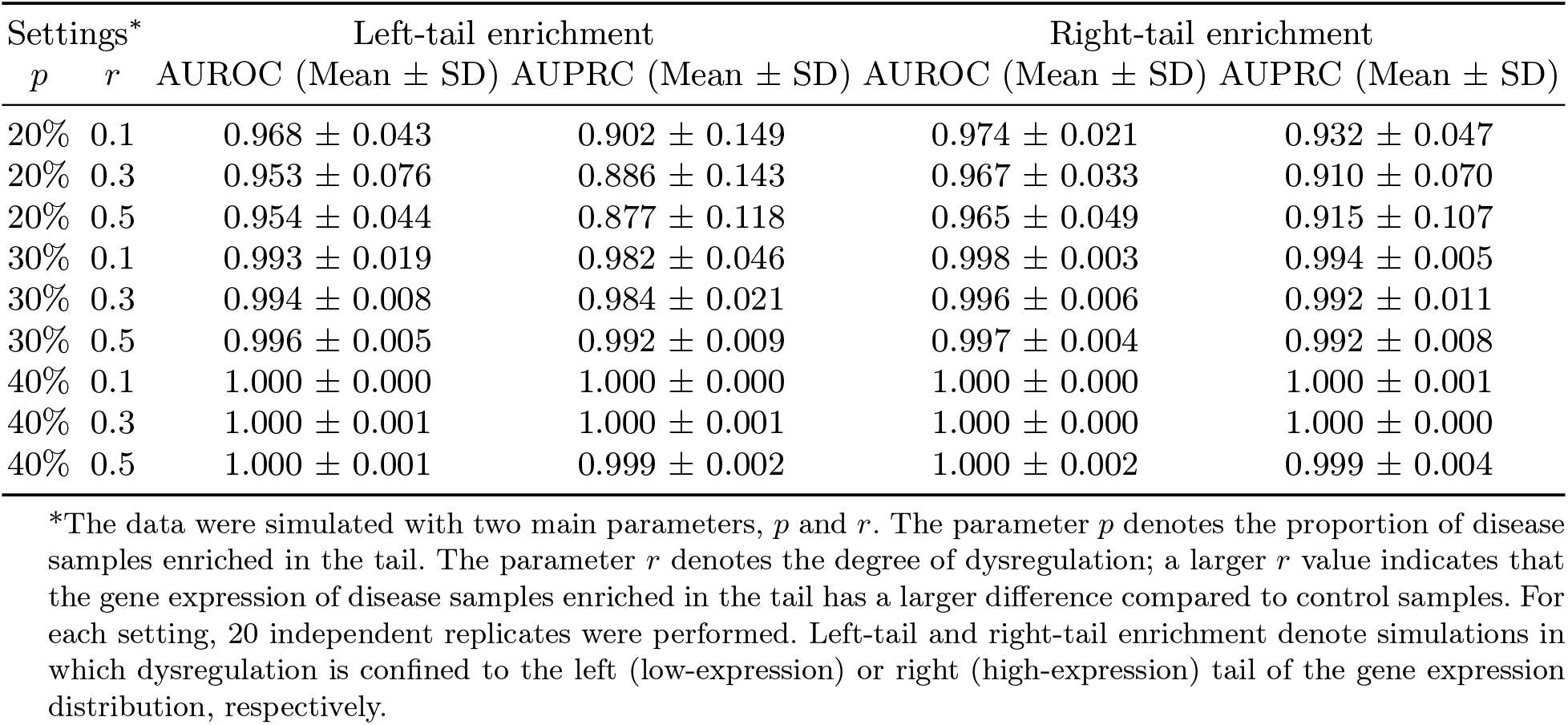
Results of TailEnrich in simulation data with different parameter settings.

| Settings<sup>*</sup> |  | Left-tail enrichment |  | Right-tail enrichment |  |
|---|---|---|---|---|---|
| *p* | *r* | AUROC (Mean ± SD) | AUPRC (Mean ± SD) | AUROC (Mean ± SD) | AUPRC (Mean ± SD) |
| 20% | 0.1 | 0.968 ± 0.043 | 0.902 ± 0.149 | 0.974 ± 0.021 | 0.932 ± 0.047 |
| 20% | 0.3 | 0.953 ± 0.076 | 0.886 ± 0.143 | 0.967 ± 0.033 | 0.910 ± 0.070 |
| 20% | 0.5 | 0.954 ± 0.044 | 0.877 ± 0.118 | 0.965 ± 0.049 | 0.915 ± 0.107 |
| 30% | 0.1 | 0.993 ± 0.019 | 0.982 ± 0.046 | 0.998 ± 0.003 | 0.994 ± 0.005 |
| 30% | 0.3 | 0.994 ± 0.008 | 0.984 ± 0.021 | 0.996 ± 0.006 | 0.992 ± 0.011 |
| 30% | 0.5 | 0.996 ± 0.005 | 0.992 ± 0.009 | 0.997 ± 0.004 | 0.992 ± 0.008 |
| 40% | 0.1 | 1.000 ± 0.000 | 1.000 ± 0.000 | 1.000 ± 0.000 | 1.000 ± 0.001 |
| 40% | 0.3 | 1.000 ± 0.001 | 1.000 ± 0.001 | 1.000 ± 0.000 | 1.000 ± 0.000 |
| 40% | 0.5 | 1.000 ± 0.001 | 0.999 ± 0.002 | 1.000 ± 0.002 | 0.999 ± 0.004 |

\*The data were simulated with two main parameters, *p* and *r*. The parameter *p* denotes the proportion of disease samples enriched in the tail. The parameter *r* denotes the degree of dysregulation; a larger *r* value indicates that the gene expression of disease samples enriched in the tail has a larger difference compared to control samples. For each setting, 20 independent replicates were performed. Left-tail and right-tail enrichment denote simulations in which dysregulation is confined to the left (low-expression) or right (high-expression) tail of the gene expression distribution, respectively.

**Fig 2.**
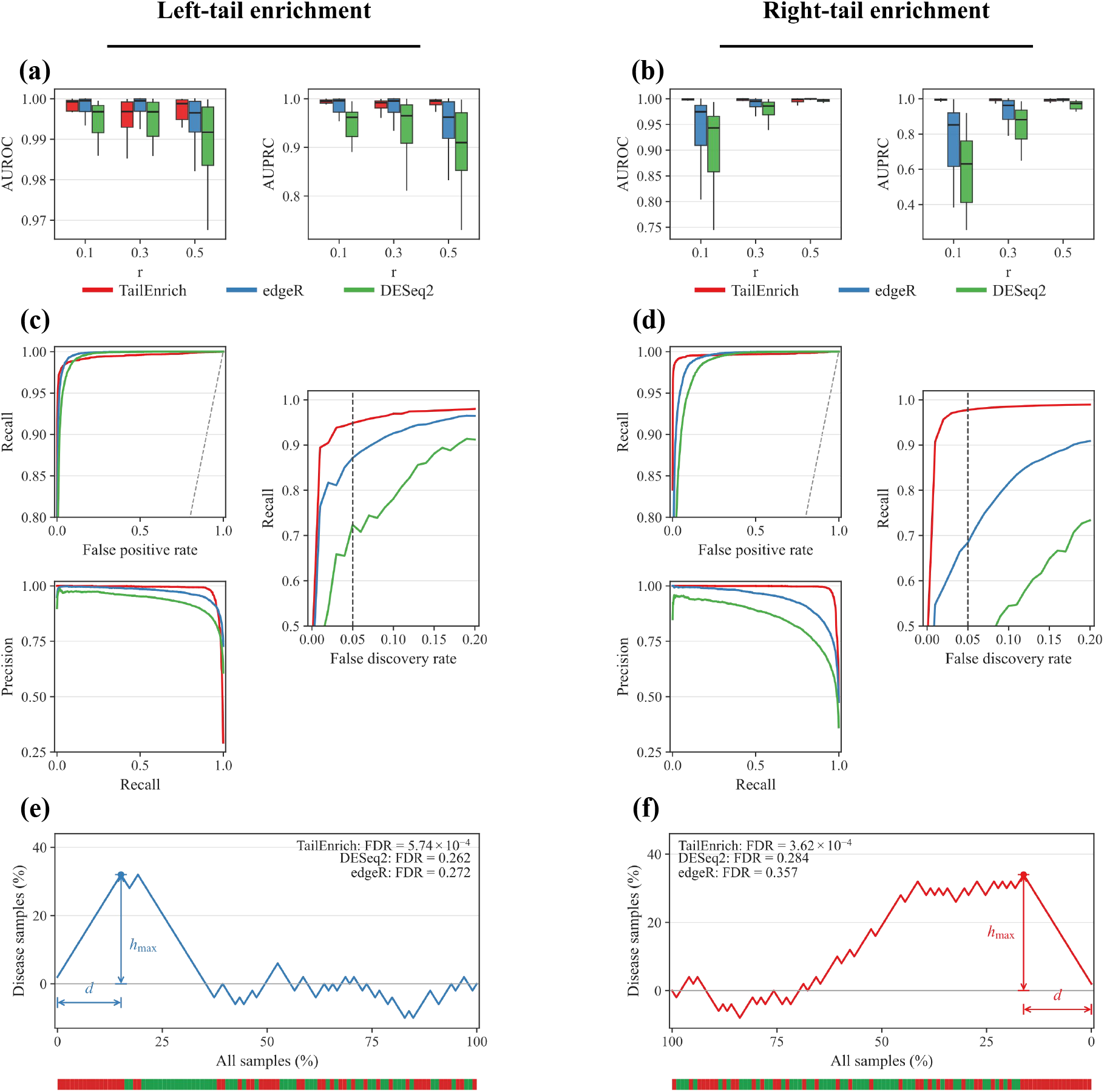
Performance comparison of TailEnrich, DESeq2, and edgeR in distinguishing dysregulated from non-dysregu lated genes on simulated datasets. **(a**,**b)** AUROC and AUPRC comparisons under left-tail and right-tail enrichment, respectively, with *p* = 30% and *r* = 0.1, 0.3, and 0.5. For each parameter setting, 20 replicate experiments were run. **(c**,**d)** ROC, precision–recall (PR), and FDR–Recall curves under left-tail and right-tail enrichment, respectively, with *p* = 30% and *r* = 0.3. ROC and PR curves were averaged over 20 independent replicates, and FDR–Recall curves show mean Recall across 20 independent replicates in the low-FDR region. Dashed vertical lines in the FDR–Recall plots indicate *FDR* = 0.05. **(e**,**f)** The pattern of percentage of enriched disease samples for two representative genes with left-tail and right-tail enrichment, respectively. The color strip below each curve indicates the sample group at each ranked position; green and red denote control and disease samples, respectively.

To illustrate changes in the Percentage of Enriched disease samples across ranked positions, we selected one gene from the left-tail simulation and one gene from the right-tail simulation with *p* = 30% and *r* = 0.3 (Fig. 2e,f). For the gene from the left-tail simulation, *h*max occurred near the low-expression tail. TailEnrich identified this gene as significant (FDR = 5.74 *×* 10^*−*4^). DESeq2 (FDR = 0.262) and edgeR (FDR = 0.272) did not identify it as significant (Fig. 2e). For the gene from the right-tail simulation, *h*max occurred near the high-expression tail. TailEnrich identified this gene as significant (FDR = 3.62 *×* 10^*−*4^). DESeq2 (FDR = 0.284) and edgeR (FDR = 0.357) did not identify it as significant (Fig. 2f).

### 3.2 TailEnrich identifies AD-associated genes which cannot be uncovered by DESeq2 and edgeR

We compared TailEnrich with DESeq2 and edgeR on two public AD RNA-seq datasets: MayoRNAseq tem poral cortex (MayoRNAseq-TCX; 69 controls and 80 AD samples) tissue [9] and parahippocampal gyrus samples from Brodmann area 36 in the Mount Sinai Brain Bank (MSBB-BM36; 38 controls and 58 AD sam ples) [10]. Their Synapse study IDs are syn5550404 (MayoRNAseq) and syn3159438 (MSBB). Isoform-level TPM values and read counts were obtained from raw FASTQ reads using Salmon [11]. Within each dataset, isoforms with TPM *≥* 1 in at least 50% of all samples were retained. The retained isoform-level TPM val ues and read counts were then aggregated to compute the corresponding gene-level expression matrices for subsequent analyses.

Covariates were adjusted for each dataset. For MayoRNAseq-TCX, the covariates were sex, age at death, APOE genotype, RNA integrity number, and sample source. For MSBB-BM36, the covariates were sex, race, age at death, APOE genotype, and postmortem interval. Before running TailEnrich, covariate effects were removed from the gene-level TPM matrix by linear regression. The adjusted TPM matrix and disease-group labels were then used as inputs to TailEnrich. In contrast, DESeq2 and edgeR used the unadjusted gene-level count matrix and directly incorporated the same covariates into their respective design matrices. The detailed procedures for expression filtering and covariate adjustment are provided in Supplementary Note S2.

#### TailEnrich-specific genes show significant association with Alzheimer’s disease

We benchmarked TailEnrich against DESeq2 (version 1.42.0) and edgeR (version 4.0.16) in detecting differential genes across the two AD datasets (Fig. 3a). Significance was defined as *FDR <* 0.05 and |log2 FC| *≥* log2(1.5). The calculation of log2 FC for TailEnrich is described in Supplementary Note S3. TailEnrich identified 1,533 and 2,007 significant genes in MayoRNAseq-TCX and MSBB-BM36, respectively, compared with 1,397 and 807 for DESeq2 and 1,392 and 782 for edgeR. The overlapping genes detected by these three methods are shown in Fig. 3a, with 790 and 543 genes identified by all three methods. Genes detected by TailEnrich but by neither DESeq2 nor edgeR in the same dataset were defined as TailEnrich-specific genes. This analysis yielded 713 and 1,444 TailEnrich-specific genes in MayoRNAseq-TCX and MSBB-BM36, respectively. Two example genes illustrate differences among the three methods (Fig. 3b). For example, in MSBB-BM36, *RGS2* was identified as significant by TailEnrich (*FDR* = 0.016), but not by DESeq2 (*FDR* = 0.754) or edgeR (*FDR* = 0.771); and in MayoRNAseq-TCX, *IL33* was identified as significant by TailEnrich (*FDR* = 1.6 *×* 10^*−*3^), but not by DESeq2 (*FDR* = 0.119) or edgeR (*FDR* = 0.110).

**Fig 3.**
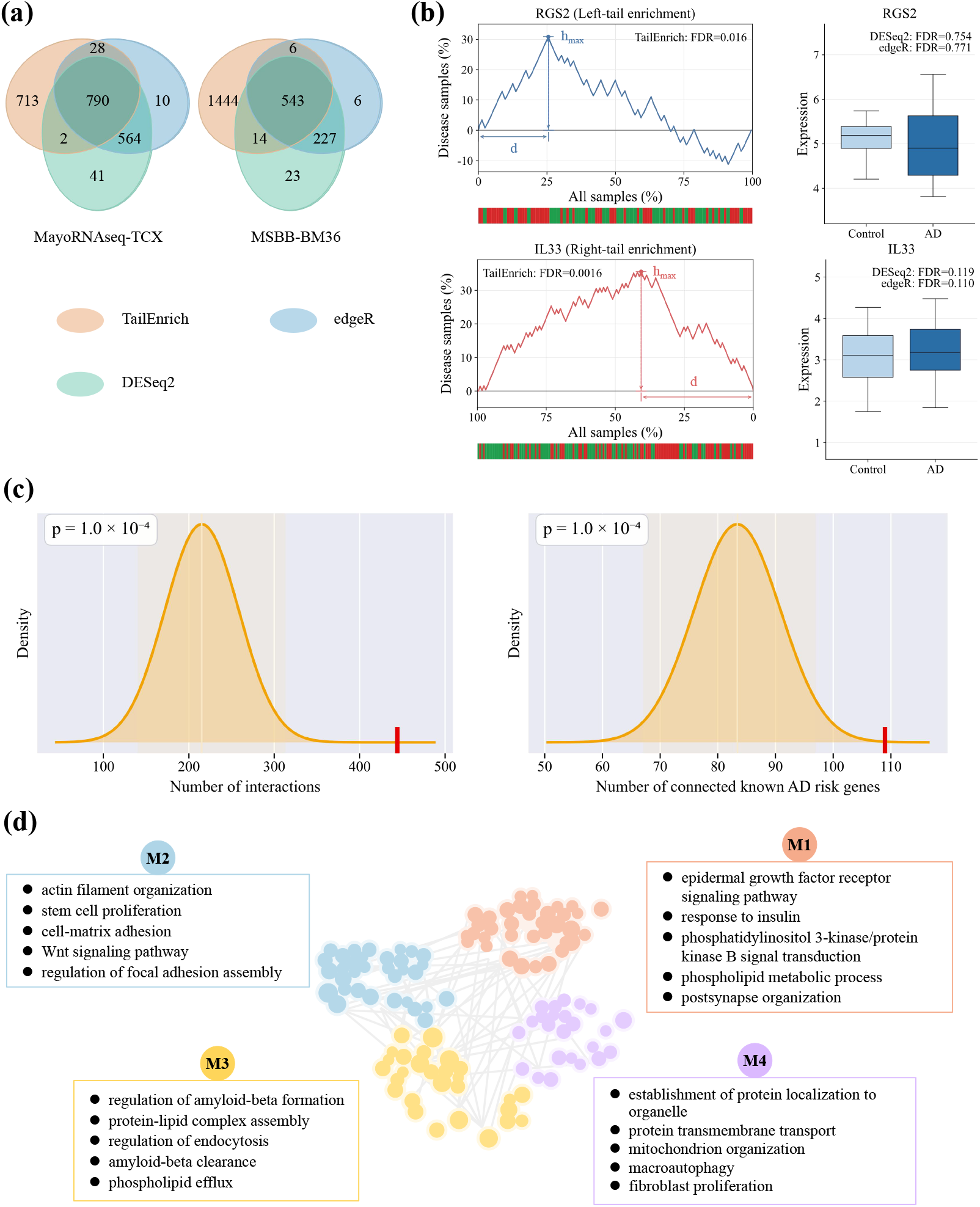
Analysis of TailEnrich-specific genes in Alzheimer’s disease cohorts. **(a)** Overlap of significant genes (*FDR <* 0.05, |log2 FC| *≥* log2(1.5)) identified by TailEnrich, DESeq2, and edgeR in MayoRNAseq-TCX and MSBB-BM36. **(b)** Tail-enrichment profiles and between-group expression comparisons for two example TailEnrich-specific genes, *IL33* in MayoRNAseq-TCX and *RGS2* in MSBB-BM36. The color strip below each enrichment curve indicates the sample group at each ranked position; green and red denote control and AD samples, respectively. **(c)** The top 200 TailEnrich-specific genes show significant interactions with known AD risk genes; this analysis was performed using the AlzCode webserver. **(d)** The four retained network modules (M1, M2, M3, and M4), each containing at least 15 top 200 genes, are annotated with representative GO Biological Process terms.

The top 200 TailEnrich-specific genes displayed significant network interactions with known AD risk genes. We collected a set of 239 known AD risk genes (Supplementary Table S5) from 33 GWAS studies in the NHGRI–EBI GWAS Catalog [12]. Using the Gene Set Analysis tool in the AlzCode webserver [13], we assessed interactions between the top 200 TailEnrich-specific genes and the 239 known AD risk genes within the ADBrainNexus framework [14]. Of the top 200 TailEnrich-specific genes, 114 had at least one interaction. The observed number of connected known AD risk genes was 109, compared with a random range of 67–97 (*p* = 1.0 *×* 10^*−*4^). The observed number of interactions was 444, compared with a random range of 140–313 (*p* = 1.0 *×* 10^*−*4^; Fig. 3c).

We identified gene modules using the top 200 TailEnrich-specific genes. These genes and the known AD risk genes were combined to construct a subnetwork from ADBrainNexus by retaining edges with weights *>* 0.75. The Louvain algorithm implemented in NetworkX (version 3.1) [15] was used to partition the subnetwork. This partitioning resulted in 12 modules, of which four modules containing at least 15 top-ranked genes were retained for enrichment analysis (Fig. 3d). The sizes and compositions of these four modules, i.e. M1, M2, M3, and M4, are provided in Supplementary Table S6.

The network modules were enriched for AD-related biological processes (Fig. 3d). We performed Gene Ontology enrichment for each module using the enrichGO function in the clusterProfiler package (version 4.18.4) [16]. Enriched GO terms containing *≤* 500 annotated genes were retained. M1 was mainly enriched for growth factor and insulin–PI3K/AKT signaling. It was reviewed in Miao et al. work that abnormalities in insulin-like growth factor signaling and its downstream PI3K/AKT pathway occur in AD [17]. Leung et al. linked an AD risk variant to increased expression of EGFR, which encodes the epidermal growth factor receptor, and confirmed that the regulatory region containing this variant influenced EGFR expression in human microglial cells [18]. M2 was mainly enriched for cytoskeletal remodeling and cell adhesion. For example, Saleh et al. found that over expression of focal adhesion kinase, which regulates cell adhesion and cytoskeletal remodeling, increased amyloid-*β* burden, altered neuronal morphology, and impaired learning and memory in an AD mouse model [19]. M3 was mainly enriched for endocytic processes. In a previous review, Jaye et al. described clathrin-mediated endocytosis as a cellular uptake route involved in amyloid-*β* processing and clearance in AD [20]. Li et al. also described how the endocytic receptor LRP1 contributes to amyloid-*β* internalization, degradation, and clearance [21]. M4 was mainly enriched for mitochondrial quality control. Percy et al. reviewed alterations in several components of mitochondrial quality control in AD, including mitochondrial proteostasis, dynamics, and mitophagy [22]. In a recent study, Dan et al. also observed abnormal mitochondrial buildup in both postmortem human AD brains and AD model mice [23].

#### Case study of TailEnrich-specific genes

In this case study, we evaluated the associations of TailEn rich-specific genes detected in MayoRNAseq-TCX with AD-related clinical and neuropathologic traits. The 713 TailEnrich-specific genes were used as input, of which 645 had expression data in MSBB-BM36. For each gene, expression was tested for correlation with Braak stage, Clinical Dementia Rating (CDR), and CERAD scores using two-sided Pearson correlation. For each trait, the 645 *p* values were adjusted using the Benjamini–Hochberg method. A gene was considered associated with a trait when *FDR <* 0.05 and |*r*| *≥*0.30. With this criterion, 114, 139, and 188 genes were found to be associated with Braak stage, CDR, and CERAD scores, respectively. Overall, 225 genes met the criteria for at least one trait, including 82 genes that met them for all three traits. Full correlation results for these 225 genes are provided in Supplementary Table S7. The top 10 of the 82 genes, ranked in descending order by the mean of their *−* log10(*FDR*) values across the three traits, are shown in Fig. 4a.

**Fig 4.**
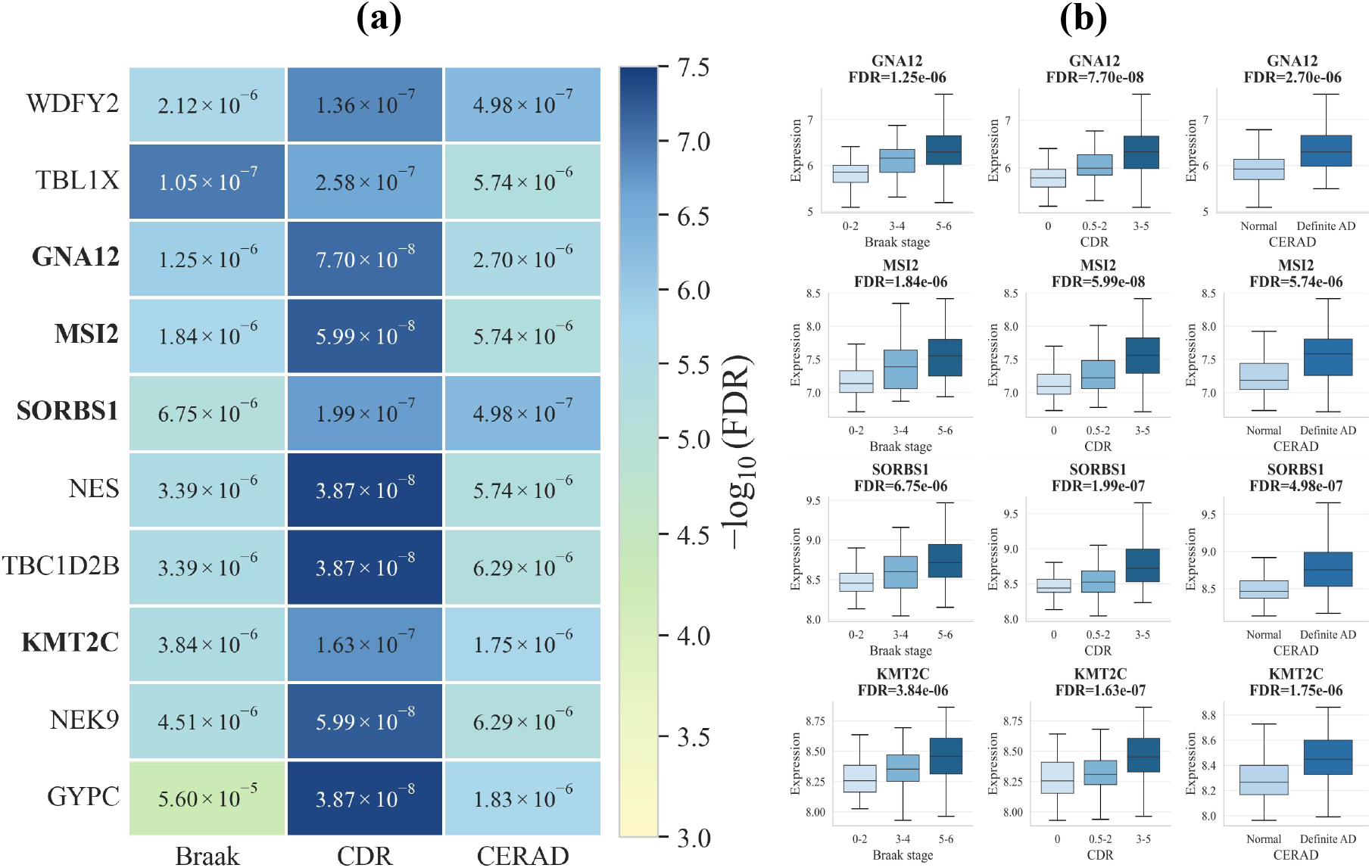
Case study of TailEnrich-specific genes in MSBB-BM36. Of the 713 TailEnrich-specific genes detected in MayoRNAseq-TCX and used as input, 645 were available for analysis in MSBB-BM36. Associations between gene expression and Braak stage, Clinical Dementia Rating (CDR), and CERAD score were assessed using two-sided Pearson correlation, with significance defined as *FDR <* 0.05 and |*r*| *≥* 0.30. **(a)** Correlation results for the top 10 of the 82 genes that met both criteria for all three traits, ranked in descending order by the mean of their *−* log10(*FDR*) values across the three traits. The text in each cell of the heatmap shows FDR values, and colors represent *−* log10(*FDR*). **(b)** Expression distributions of *GNA12, KMT2C, SORBS1*, and *MSI2* across Braak stage, CDR, and CERAD in MSBB-BM36.

Among these genes, *GNA12, MSI2, SORBS1*, and *KMT2C* have been linked to AD in previous studies. Their expression increased consistently with greater clinical and neuropathologic severity (Fig. 4b). Genetic variants near *GNA12* were associated with cerebrospinal fluid total tau and phosphorylated tau 181 levels in a genome-wide association meta-analysis [24]. At the protein level, increased MSI2 signals and an association between MSI2 and tau aggregates were observed in AD brain tissue. Tau also promoted MSI2 aggregation in vitro [25]. Li et al. reported increased *SORBS1* expression in astrocytes from AD brains and found that *SORBS1* promoted astrocyte senescence in vitro [26]. At the transcript level, *KMT2C* mRNA was significantly increased in postmortem prefrontal cortex from patients with AD compared with controls [27].

## 4 Conclusion

TailEnrich is a computational method for identifying differential genes with disease sample enrichment at the tails of their expression distributions. Conventional differential analysis methods, such as DESeq2 and edgeR (v4), mainly test mean expression differences between disease and control groups and may miss genes that are dysregulated only in a subset of disease samples. TailEnrich instead tests whether disease samples are enriched at the low- or high-expression tail of each gene. This design enables TailEnrich to identify genes with expression dysregulation that may be missed by mean-shift based methods.

The simulation and real-world AD RNA-seq data analyses support the use of TailEnrich as a comple ment to conventional differential analysis methods. In the simulation study, TailEnrich better distinguished genes dysregulated in a subset of disease samples than DESeq2 and edgeR. Across the two AD RNA-seq datasets, TailEnrich identified genes that were not detected by either DESeq2 or edgeR. TailEnrich-specific genes showed significant network-level connectivity to known AD risk genes and were enriched for AD-re lated biological processes. The case study further showed associations between TailEnrich-specific genes and Alzheimer’s clinical and neuropathologic traits, including Braak stage, CDR, and CERAD scores.

This work has several limitations that remain to be addressed in future research. First, TailEnrich is currently applied to bulk RNA-seq and may therefore miss genes that show significant dysregulation in specific cell types. Future work will extend TailEnrich to address the high sparsity and noise characteristic of single-cell RNA-sequencing data, thereby enabling the identification of genes dysregulated in specific cell types. Second, in the present study, we used Alzheimer’s disease gene expression data as a proof-of-concept application. The method could also be adapted to genomic variant and proteomic data to identify disease-as sociated variants and proteins enriched in patient subgroups. These extensions will broaden the applicability of TailEnrich to complex disease studies and facilitate the discovery of novel disease-associated molecular signals.

## Acknowledgments

This work was in part supported by Brain Science and Brain-like Intelligence Technology-Na tional Science and Technology Major Project (No. 2022ZD0213700), National Natural Science Foundation of China (No. 62473382,62302426), Hunan Provincial Natural Science Foundation of China (2025JJ20068). The results pub lished here are in part based on data obtained from the AMP-AD Knowledge Portal (https://adknowledgeportal.synapse.org/). The MayoRNAseq data are available from the following sources: The Mayo Clinic Alzheimer’s Disease Ge netic Studies, led by Dr Nilufer Ertekin-Taner and Dr Steven G. Younkin, Mayo Clinic, Jacksonville, FL using samples from the Mayo Clinic Study of Aging, the Mayo Clinic Alzheimer’s Disease Research Center and the Mayo Clinic Brain Bank. Data collection was supported through funding by NIA grants P50 AG016574, R01 AG032990, U01 AG046139, R01 AG018023, U01 AG006576, U01 AG006786, R01 AG025711, R01 AG017216, R01 AG003949, NINDS grant R01 NS080820, CurePSP Foundation and support from Mayo Foundation. Study data includes samples col lected through the Sun Health Research Institute Brain and Body Donation Program of Sun City, Arizona. The Brain and Body Donation Program is supported by the National Institute of Neurological Disorders and Stroke (U24 NS072026 National Brain and Tissue Resource for Parkinson’s Disease and Related Disorders), the National Insti tute on Aging (P30 AG19610 Arizona Alzheimer’s Disease Core Center), the Arizona Department of Health Services (contract 211002, Arizona Alzheimer’s Research Center), the Arizona Biomedical Research Commission (contracts 4001, 0011, 05-901 and 1001 to the Arizona Parkinson’s Disease Consortium) and the Michael J. Fox Foundation for Parkinson’s Research. The MSBB data are generated from postmortem brain tissue collected through the Mount Sinai VA Medical Center Brain Bank and are provided by Dr Eric Schadt from Mount Sinai School of Medicine.

## Reference

1. Tijms, B. M. et al. Cerebrospinal fluid proteomics in patients with Alzheimer’s disease reveals five molecular subtypes with distinct genetic risk profiles. Nature Aging 4, 33–47 (2024).

2. Malliou, G. et al. Distinct CSF lipidomic profiles are associated with five proteomic subtypes in patients with Alzheimer’s disease. Molecular Neurodegeneration Advances 2, 11 (2026).

3. Zolotareva, O. et al. Identification of differentially expressed gene modules in heterogeneous diseases.Bioinformatics 37, 1691–1698 (2021).

4. Love, M. I., Huber, W. & Anders, S. Moderated estimation of fold change and dispersion for RNA-seq data with DESeq2. Genome Biology 15, 550 (2014).

5. Chen, Y. et al. edgeR v4: powerful differential analysis of sequencing data with expanded functionality and improved support for small counts and larger datasets. Nucleic Acids Research 53, gkaf018 (2025).

6. Benjamini, Y. & Hochberg, Y. Controlling the false discovery rate: a practical and powerful approach to multiple testing. Journal of the Royal Statistical Society: Series B (Methodological) 57, 289–300 (1995).

7. Soneson, C. compcodeR—an R package for benchmarking differential expression methods for RNA-seq data. Bioinformatics 30, 2517–2518 (2014).

8. Wu, J. et al. Revisiting differential expression analysis: An updated six-dimensional comparative study.PLOS ONE 21, e0344709 (2026).

9. Allen, M. et al. Human whole genome genotype and transcriptome data for Alzheimer’s and other neurodegenerative diseases. Scientific Data 3, 160089 (2016).

10. Wang, M. et al. The Mount Sinai cohort of large-scale genomic, transcriptomic and proteomic data in Alzheimer’s disease. Scientific Data 5, 180185 (2018).

11. Patro, R. et al. Salmon provides fast and bias-aware quantification of transcript expression. Nature Methods 14, 417–419 (2017).

12. Cerezo, M. et al. The NHGRI–EBI GWAS Catalog: standards for reusability, sustainability and diver sity. Nucleic Acids Research 53, D998–D1005 (2025).

13. Lin, C.-X. et al. AlzCode: a platform for multiview analysis of genes related to Alzheimer’s disease.Bioinformatics 38, 2030–2032 (2022).

14. Lin, C.-X. et al. An integrated brain-specific network identifies genes associated with neuropathologic and clinical traits of Alzheimer’s disease. Briefings in Bioinformatics 23, bbab522 (2022).

15. Hagberg, A. A., Schult, D. A. & Swart, P. J. Exploring network structure, dynamics, and function using NetworkX. In Proceedings of the 7th Python in Science Conference (eds Varoquaux, G., Vaught, T. & Millman, J.) 11–15 (Pasadena, CA, USA, 2008).

16. Wu, T. et al. clusterProfiler 4.0: A universal enrichment tool for interpreting omics data. The Innovation2, 100141 (2021).

17. Miao, J. et al. Insulin-Like Growth Factor Signaling in Alzheimer’s Disease: Pathophysiology and Ther apeutic Strategies. Molecular Neurobiology 62, 3195–3225 (2025).

18. Leung, Y. Y. et al. Integrated genomic analysis and CRISPRi implicates EGFR in Alzheimer’s disease risk. npj Dementia 1, 42 (2025).

19. Saleh, B. et al. FAK-Mediated Signaling Controls Amyloid Beta Overload, Learning and Memory Deficits in a Mouse Model of Alzheimer’s Disease. International Journal of Molecular Sciences 23, 9055 (2022).

20. Jaye, S., Sandau, U. S. & Saugstad, J. A. Clathrin mediated endocytosis in Alzheimer’s disease: cell type specific involvement in amyloid beta pathology. Frontiers in Aging Neuroscience 16, 1378576 (2024).

21. Li, W. et al. The LDL Receptor-Related Protein 1: Mechanisms and roles in promoting Aβ efflux transporter in Alzheimer’s disease. Biochemical Pharmacology 231, 116643 (2025).

22. McGill Percy, K. C., Liu, Z. & Qi, X. Mitochondrial dysfunction in Alzheimer’s disease: Guiding the path to targeted therapies. Neurotherapeutics 22, e00525 (2025).

23. Dan, X. et al. Mitochondrial accumulation and lysosomal dysfunction result in mitochondrial plaques in Alzheimer’s disease. Nature Neuroscience 29, 2469–2479 (2026).

24. Timsina, J. et al. GWAS meta-analysis of cerebrospinal fluid Alzheimer’s biomarkers reveals loci regu lating lipids, brain volume and autophagy. Nature Communications 17, 7385 (2026).

25. Montalbano, M. et al. RNA-binding proteins Musashi and tau soluble aggregates initiate nuclear dys function. Nature Communications 11, 4305 (2020).

26. Li, Y. & He, X. Single-cell Transcriptomics Reveals that the SORBS1/FBXO22/BAG3 Axis Drives Astrocyte Senescence via Calcium Signaling and Affects Alzheimer’s Disease-Related Neuronal Damage. NeuroMolecular Medicine 28, 37 (2026).

27. Cao, Q. et al. Targeting histone K4 trimethylation for treatment of cognitive and synaptic deficits in mouse models of Alzheimer’s disease. Science Advances 6, eabc8096 (2020).

