## Supplementary Materials for "TailEnrich enables discovery of dysregulated genes through disease sample enrichment testing"

### TailEnrich Supplementary Materials

List of supplementary content:

| Item | Description |
|---|---|
| Supplementary Note S1 | Simulation data generation. |
| Supplementary Note S2 | Expression processing and covariate adjustment of AD RNA-seq datasets. |
| Supplementary Note S3 | Tail-specific log2FC calculation. |
| Supplementary Figure S1 | Examples of genes with completely separated expression levels between disease and control groups. |
| Supplementary Figure S2 | Performance on simulated datasets for 5,000 genes under left-tail enrichment. |
| Supplementary Figure S3 | Performance on simulated datasets for 5,000 genes under right-tail enrichment. |
| Supplementary Figure S4 | Performance on simulated datasets for 10,000 genes under left-tail enrichment. |
| Supplementary Figure S5 | Performance on simulated datasets for 10,000 genes under right-tail enrichment. |
| Supplementary Figure S6 | Recall at low FDR on simulated datasets for 5,000 genes under left-tail enrichment. |
| Supplementary Figure S7 | Recall at low FDR on simulated datasets for 5,000 genes under right-tail enrichment. |
| Supplementary Figure S8 | Recall at low FDR on simulated datasets for 10,000 genes under left-tail enrichment. |
| Supplementary Figure S9 | Recall at low FDR on simulated datasets for 10,000 genes under right-tail enrichment. |
| Supplementary Table S1 | Simulation parameters. |
| Supplementary Table S2 | Performance of TailEnrich across all simulation settings. |
| Supplementary Table S3 | Performance of edgeR across all simulation settings. |
| Supplementary Table S4 | Performance of DESeq2 across all simulation settings. |
| Supplementary Table S5 | The 239 known AD risk genes collected from 33 GWAS studies in the NHGRI-EBI GWAS Catalog. |
| Supplementary Table S6 | Sizes and compositions of ADBrainNexus network modules. |
| Supplementary Table S7 | The expression-trait correlation results on the MSBB-BM36 dataset. |

### Supplementary Note S1. Simulation data generation

The simulated datasets were generated in two stages. First, a baseline RNA-seq count matrix without differential expression was generated using the generateSyntheticData function in the compcodeR package [1]. The primary simulation scenario contained 5,000 genes and 100 samples, including 50 disease and 50 control samples. An additional scenario containing 10,000 genes was generated using the same procedure for supplementary evaluation. No differentially expressed genes were introduced during baseline generation (n.diffexp = 0).

Second, tail enrichment was introduced into 10% of the baseline genes. In both left-tail and right-tail enrichment, two parameters, p and r, were used to control the simulated data. The parameter p specified the proportion of disease samples enriched at the tail of the gene expression distribution. The parameter r denoted the degree of dysregulation. A larger r value indicated a larger expression difference between the tail-enriched disease samples and the corresponding control-group boundary. In the simulations, p was set to 20%, 30%, or 40%, and r was set to 0.1, 0.3, or 0.5. For each setting, 20 independent replicates were generated using reproducible random seeds.

For each gene selected for tail-enrichment simulation, a proportion p of disease samples was selected for modification. Let n denote the number of disease samples. The number of disease samples selected for modification, k, was calculated as:

$$k=np.$$

For left-tail enrichment, the k disease samples with the lowest expression values were modified. Let b denote the minimum expression value among control samples. For each selected gene, let x denote the highest expression value among the k selected disease samples. The selected disease sample expression values were first shifted so that their upper boundary was aligned with b. The shift was defined as:

$$s=b-x.$$

Each selected disease sample expression value was shifted as:

$$y_i=x_i+s,$$

where $x_i$ denotes the original expression value of the selected disease sample and $y_i$ denotes the shifted value. The shifted values were then rescaled around b according to the degree of dysregulation, r.

For left-tail enrichment, the target mean of the modified disease sample expression values was defined as:

$$M_{target} = b(1-r)$$

Let $M_1$ denote the mean of the shifted disease sample expression values. The scaling factor was defined as:

$$a=\frac{M_{target}-b}{M_1-b}.$$

The final perturbed expression value was then obtained by pivot scaling around b:

$$z_i=b+a(y_i-b).$$

For right-tail enrichment, the k disease samples with the highest expression values were modified. Here, b was defined as the maximum expression value among control samples, and x was defined as the lowest expression value among the selected disease samples. The same shifting and rescaling procedure described above was then applied, with the selected disease sample boundary aligned with b and the target mean set to:

$$M_{target} = b(1+r)$$

### Supplementary Note S2. Expression processing and covariate adjustment of AD RNA-seq datasets

Two public AD RNA-seq datasets were analyzed: MayoRNAseq temporal cortex (MayoRNAseq-TCX) [2] and Mount Sinai Brain Bank Brodmann area 36 (MSBB-BM36) [3]. Raw FASTQ reads were quantified with Salmon to obtain isoform-level TPM values and read counts [4]. Gene-level expression matrices were then computed from the filtered isoform-level expression data. In each dataset, isoforms with TPM of at least 1 in at least 50% of all samples were retained. The retained isoform-level TPM values and read counts were aggregated by gene to compute the gene-level TPM and count matrices used in subsequent analyses.

Covariates were adjusted for each dataset in the subsequent analyses. For DESeq2 [5] and edgeR [6], the selected covariates were included directly in the design matrix together with the AD/control group term. For TailEnrich, covariate effects were adjusted before analysis using linear regression, following the covariate-adjustment approach described by Li et al. [7]. Briefly, TPM values were $\log_2$-transformed, and the effects of the selected covariates were regressed out while retaining the AD/control group effect. The adjusted expression values were then transformed back to the TPM scale and used as the input for TailEnrich.

### Supplementary Note S3. Tail-specific $log_2$FC calculation

TailEnrich calculates a tail-specific $log_2$FC to summarize the effect size of disease sample enrichment in the expression tail corresponding to the retained PE score. The calculation is performed separately according to whether the retained PE score represents left-tail or right-tail enrichment.

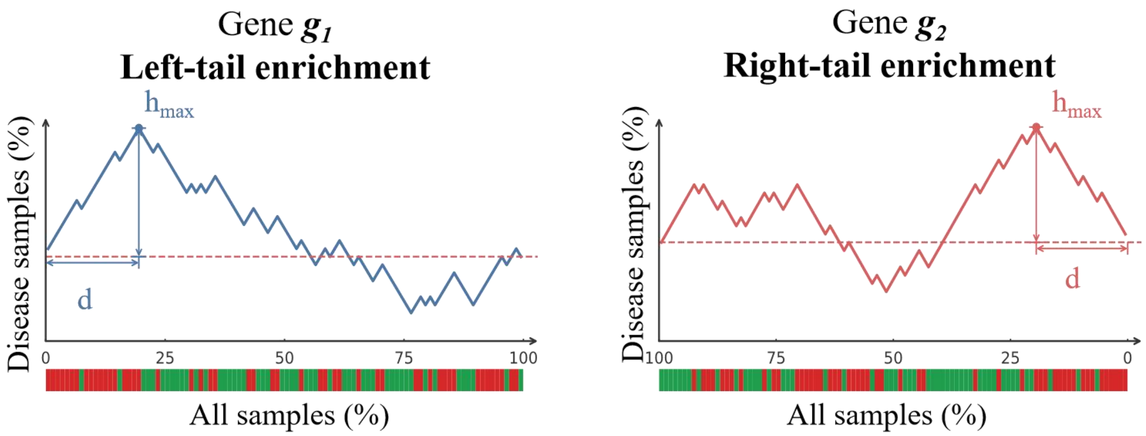


**Fig. 1. Overview of tail-specific $log_2$FC calculation.** Genes $g_1$ and g2 illustrate left-tail and right-tail enrichment, respectively. For $g_1$, the enriched region extends from $h_{max}$ to the low-expression end; for $g_2$, it extends from $h_{max}$ to the high-expression end. The mean expression of disease samples within the enriched region is used as the numerator. Starting from the corresponding expression tail, the same number of control samples are selected in the order encountered, and their mean expression is used as the denominator. The color strip below each curve indicates the sample group at each ranked position; green and red denote control and disease samples, respectively.

For left-tail enrichment, all samples were ordered from low to high expression. The position of $h_{max}$ defined the enriched region extending from the low-expression end through $h_{max}$, including the sample at $h_{max}$. Let $S_{left}$ denote the set of disease samples in this region and $m_{left}$ denote the number of these samples. Starting from the low-expression end, the complete ordered sequence was then scanned, and the first $m_{left}$ control samples encountered were selected, regardless of whether they were located inside or outside the enriched region. $D_{mean}^{left}$ was defined as the mean expression level of the disease samples in $S_{left}$, and $C_{mean}^{left}$ was defined as the mean expression level of the selected control samples. The $log_2$FC was calculated as:

$$\log_2\text{FC} = \log_2\left(\frac{D_{mean}^{left}+1}{C_{mean}^{left}+1}\right).$$

For right−tail enrichment, all samples were ordered from high to low expression. The position of $h_{max}$ defined the enriched region extending from the high−expression end through $h_{max}$, including the sample at $h_{max}$. Let $S_{right}$ denote the set of disease samples in this region and $m_{right}$ denote the number of these samples. Starting from the high−expression end, the complete ordered sequence was then scanned, and the first $m_{right}$ control samples encountered were selected, regardless of whether they were located inside or outside the enriched region. $D_{mean}^{right}$ was defined as the mean expression level of the disease samples in $S_{right}$, and $C_{mean}^{right}$ was defined as the mean expression level of the selected control samples. The $log_2$FC was calculated analogously as:

$$\log_2\text{FC} = \log_2\left(\frac{D_{mean}^{right}+1}{C_{mean}^{right}+1}\right).$$

If fewer than $m_{left}$ or $m_{right}$ control samples were available, all available control samples were used to calculate the corresponding control-group mean.

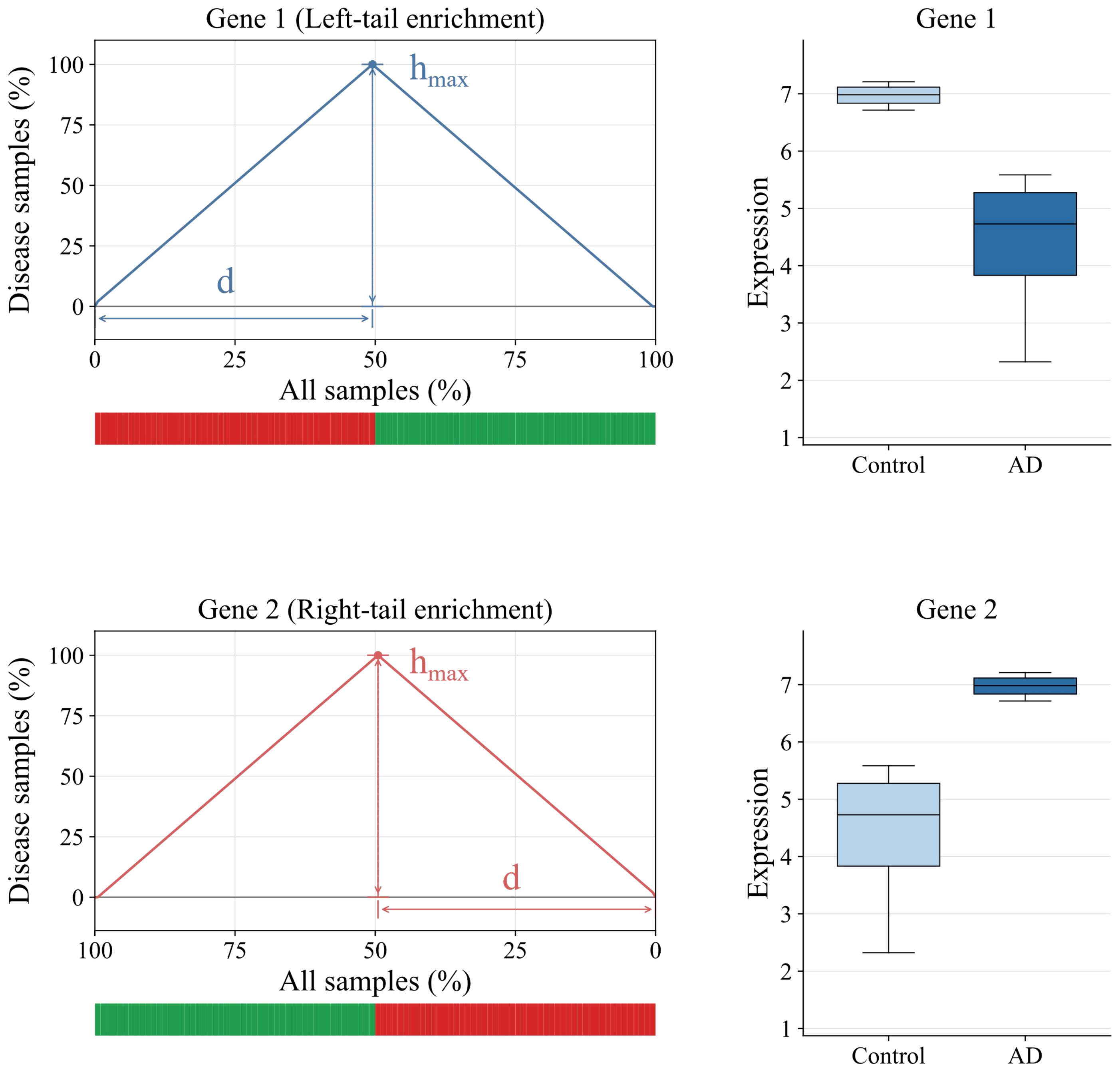


**Supplementary Figure S1. Examples of genes with completely separated expression levels between disease and control groups.** Tail-enrichment profiles and between-group expression comparisons are shown for two example genes. For these two genes, the disease and control groups are equal in size, and their expression distributions are completely separated between disease and control groups.

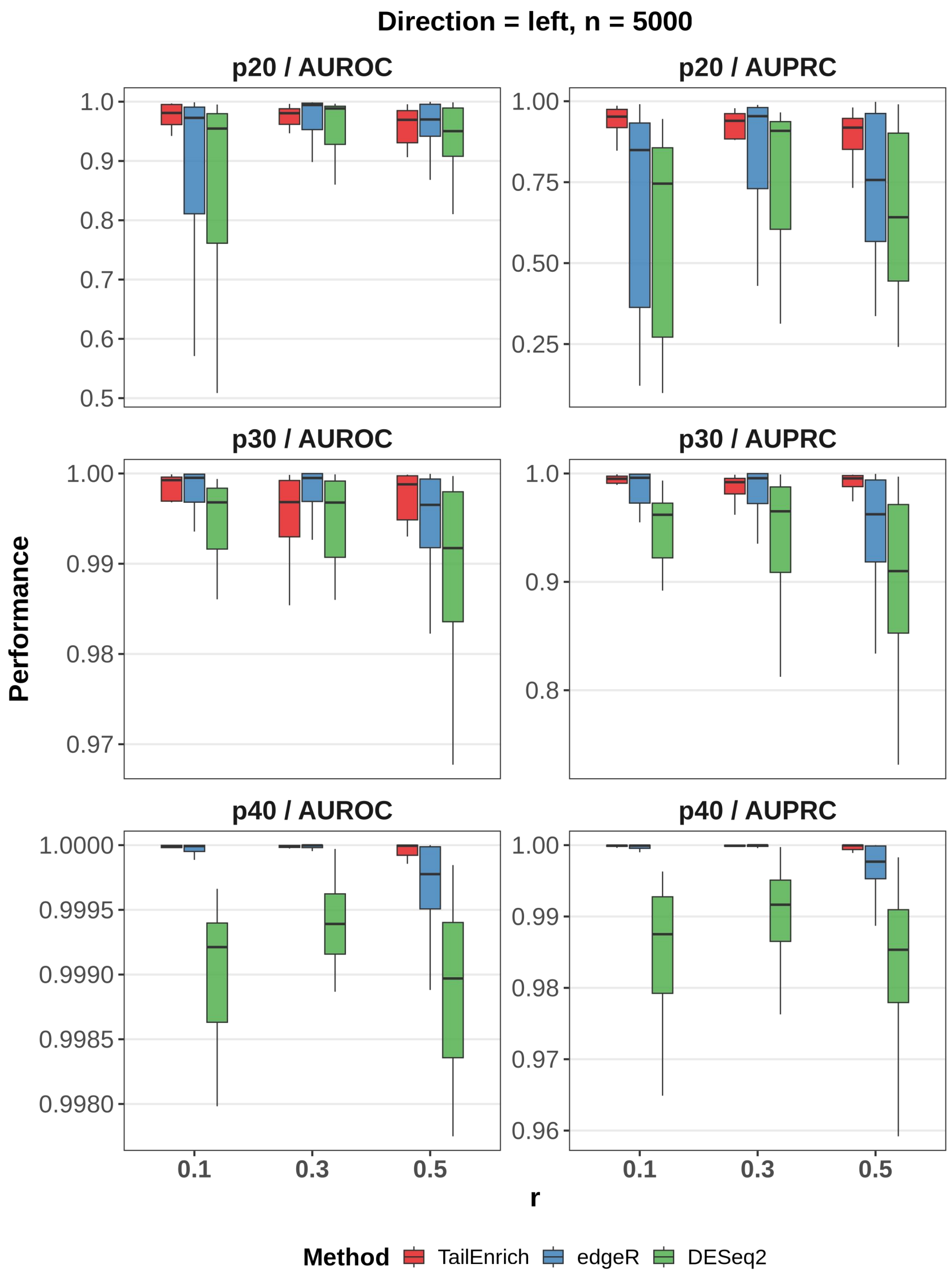


**Supplementary Figure S2. Performance on simulated datasets for 5,000 genes under left-tail enrichment.**

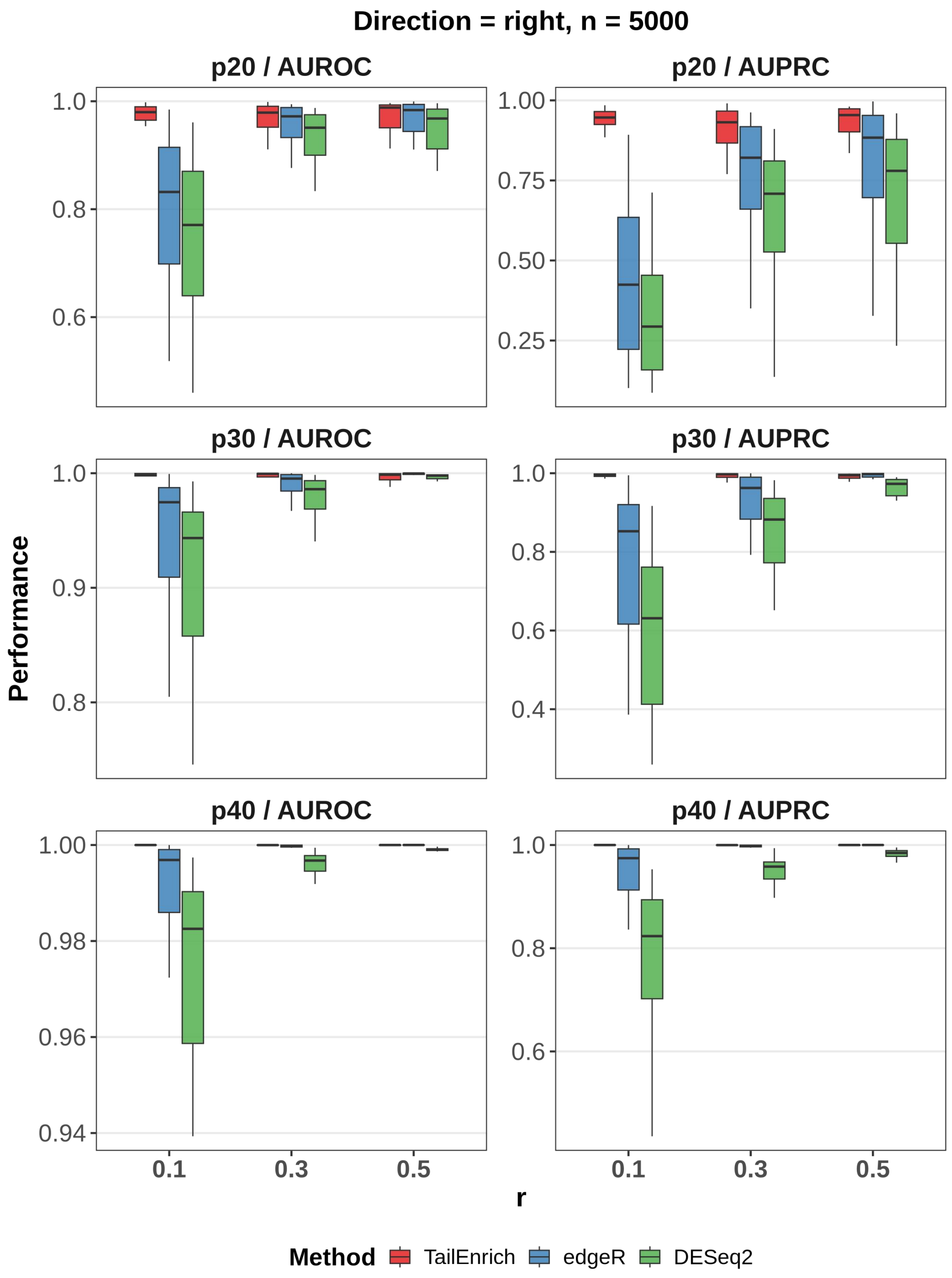


**Supplementary Figure S3. Performance on simulated datasets for 5,000 genes under right-tail enrichment.**

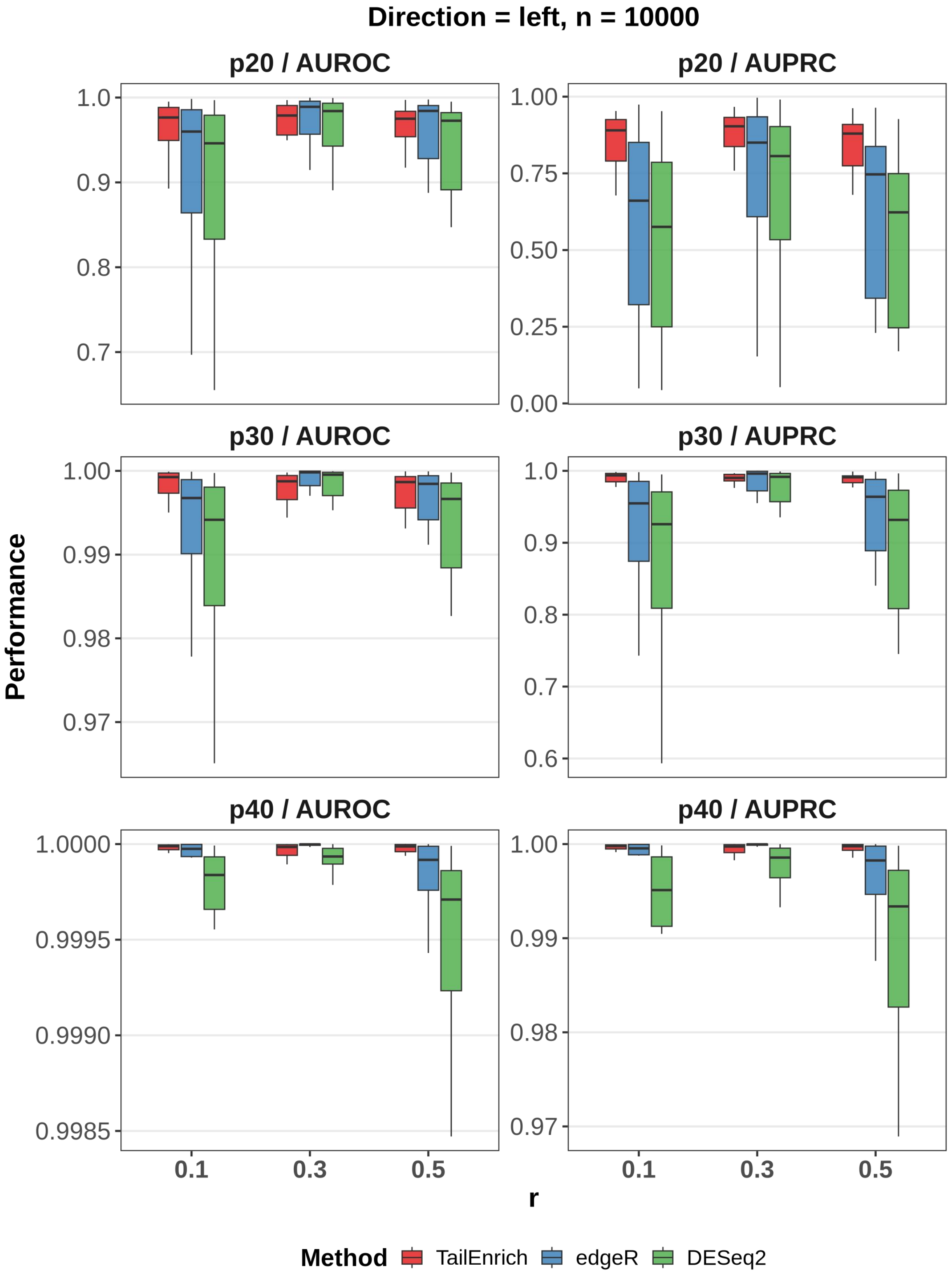


**Supplementary Figure S4. Performance on simulated datasets for 10,000 genes under left-tail enrichment.**

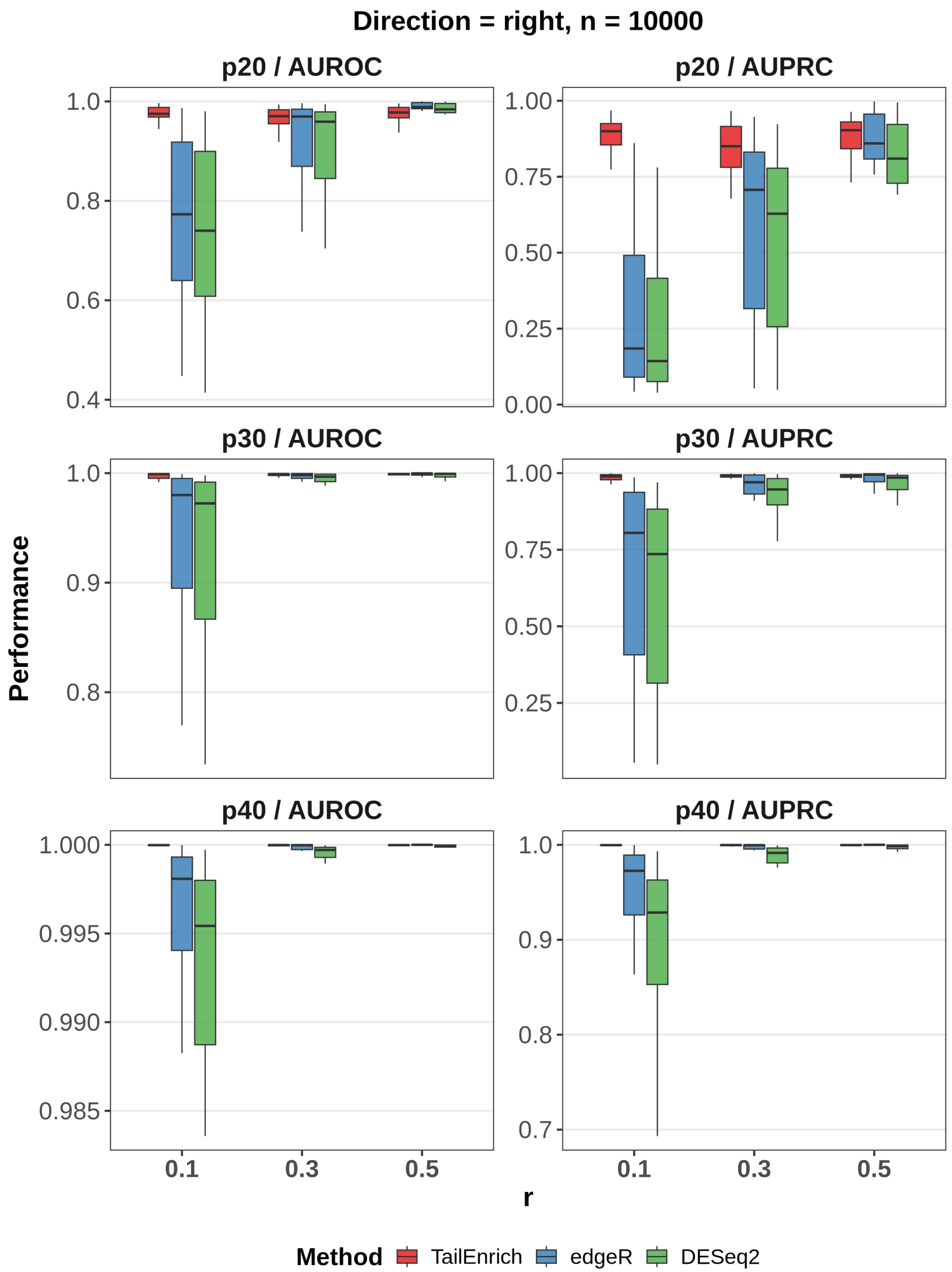


**Supplementary Figure S5. Performance on simulated datasets for 10,000 genes under right-tail enrichment.**

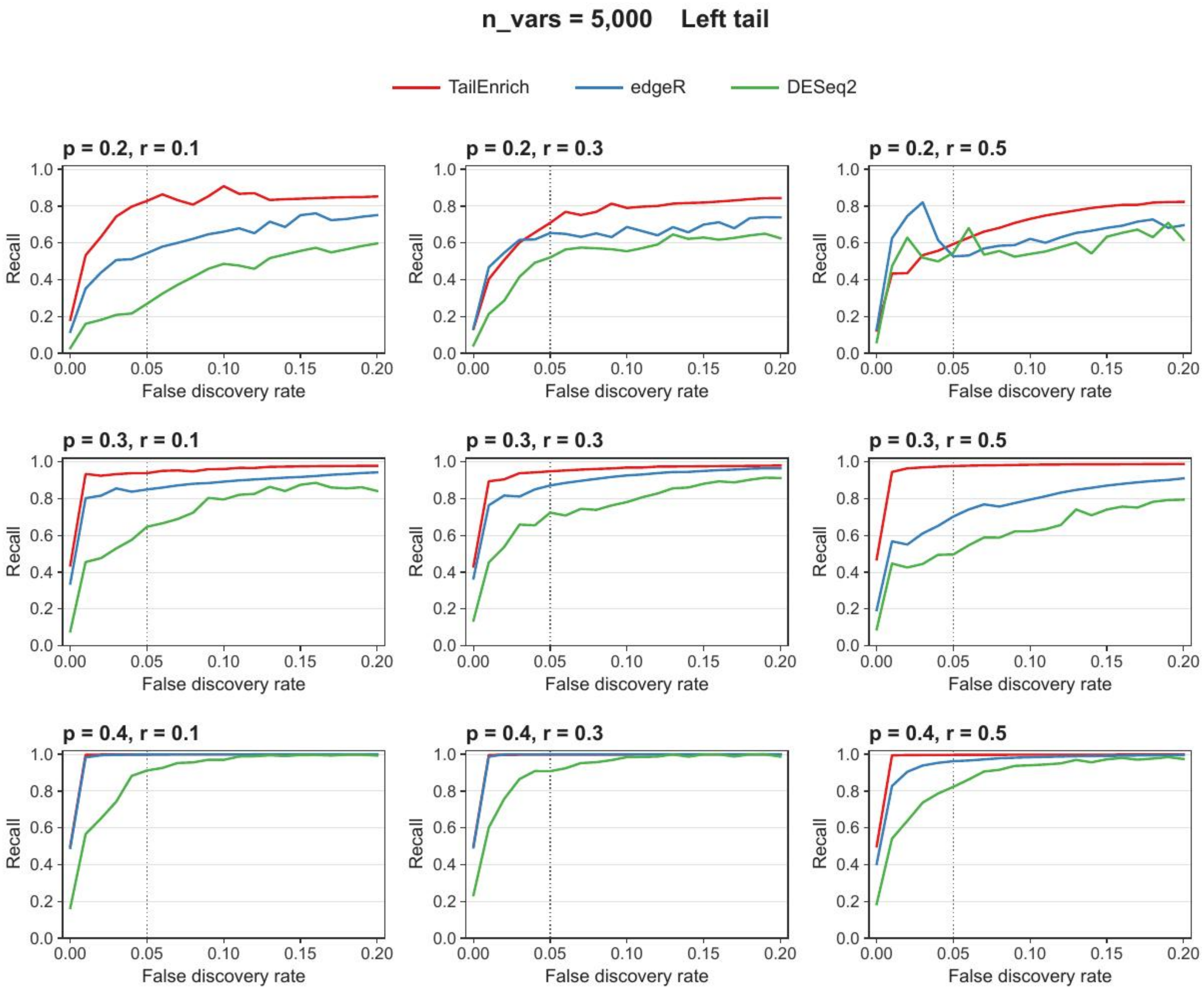


**Supplementary Figure S6. Recall at low FDR on simulated datasets for 5,000 genes under left-tail enrichment.**

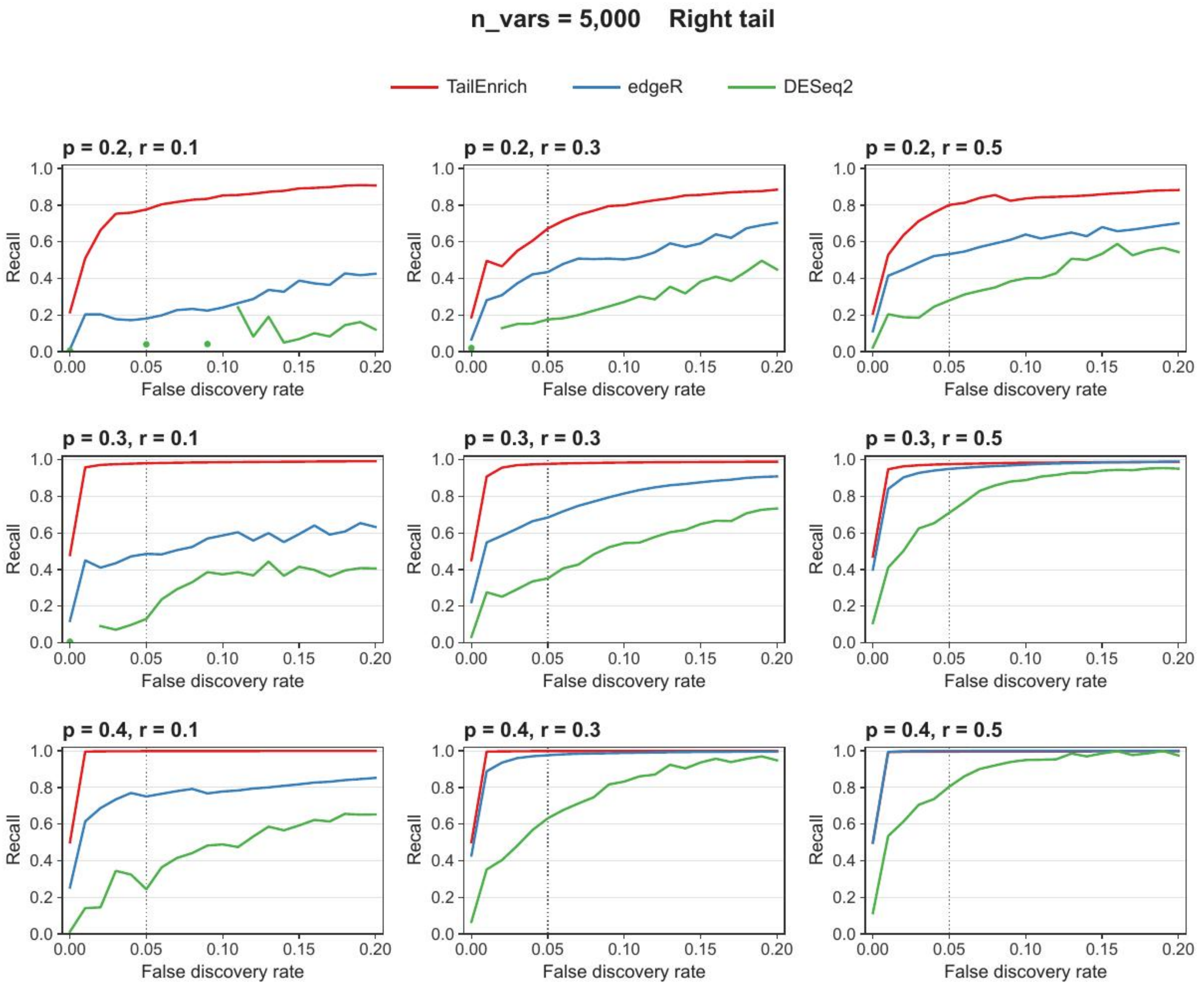


**Supplementary Figure S7. Recall at low FDR on simulated datasets for 5,000 genes under right-tail enrichment.**

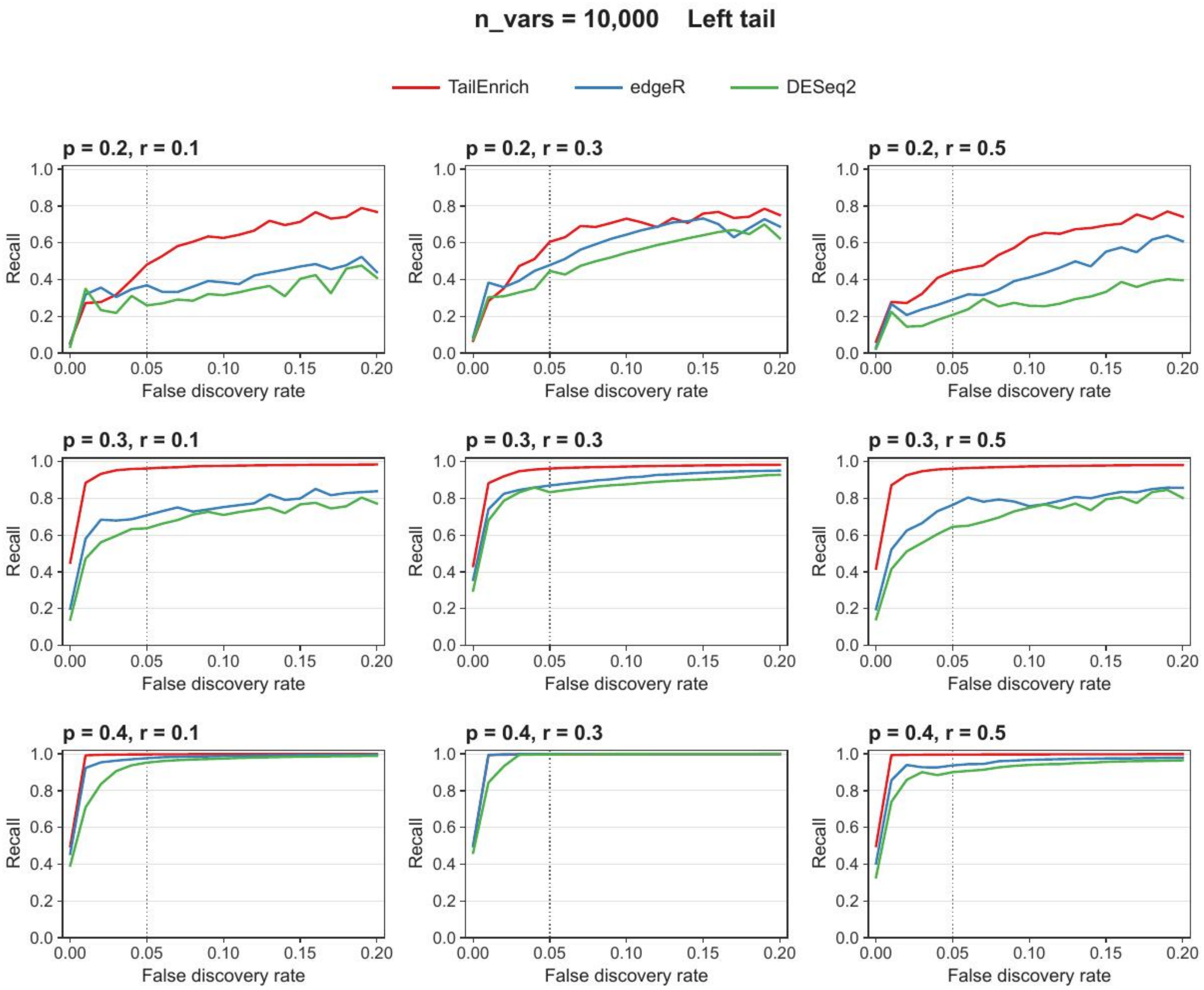


**Supplementary Figure S8. Recall at low FDR on simulated datasets for 10,000 genes under left-tail enrichment.**

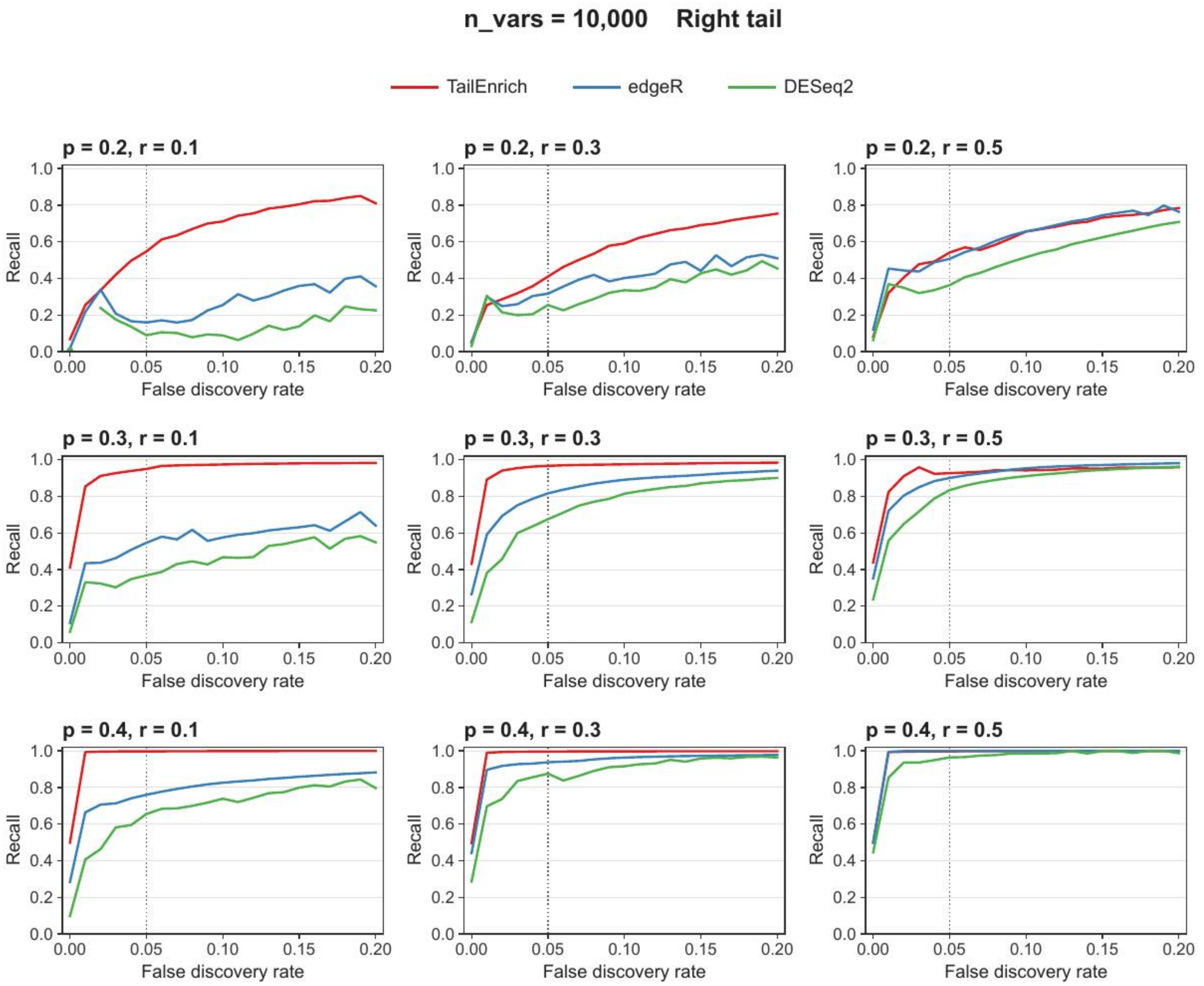


**Supplementary Figure S9. Recall at low FDR on simulated datasets for 10,000 genes under right-tail enrichment.**

### Supplementary Table S1. Simulation parameters.

| Parameter | Value | Description |
| --- | --- | --- |
| n_vars | 5000 | Number of simulated genes. |
| n_vars, supplementary scenario | 10000 | Number of simulated genes retained as an additional supplementary scenario. |
| n_tail_genes | 500 | Number of ground-truth genes. |
| n_tail_genes, supplementary scenario | 1000 | Number of ground-truth genes in the additional supplementary scenario with 10,000 simulated genes. |
| samples.per.cond | 50 | Number of samples per disease/control condition. |
| seqdepth | 5e7 | Sequencing depth parameter. |
| n.diffexp | 0 | No baseline differential expression before tail enrichment simulation. |
| tail direction | left / right | Direction of tail-specific perturbation. |
| p | 0.2, 0.3, 0.4 | Proportion of disease samples perturbed into the expression tail. |
| r | 0.1, 0.3, 0.5 | Degree of dysregulation. |
| independent replicates | 20 | Number of independent simulation replicates per setting. |
| er_cpm_cutoff | 1.0 | CPM cutoff used in expression-rate candidate filtering. |
| er_prop_r | 0.5 | Minimum expression-rate proportion used in candidate filtering. |
| b_min | 10 | Minimum eligible control-boundary value. |
| filter.threshold.total | 0 | compcodeR filtering parameter. |
| filter.threshold.mediancpm | 0 | compcodeR filtering parameter. |
| fraction.non.overdispersed | 1.0 | compcodeR dispersion-related parameter. |
| between.group.diffdisp | FALSE | No between-group differential dispersion in the baseline setting. |
| random.outlier.high.prob | 0 | Outlier-related parameter. |
| random.outlier.low.prob | 0 | Outlier-related parameter. |
| single.outlier.high.prob | 0 | Outlier-related parameter. |
| single.outlier.low.prob | 0 | Outlier-related parameter. |

### Supplementary Table S2. Performance of TailEnrich across all simulation settings.

| n_vars | p | r | Left AUROC | Left AUPRC | Right AUROC | Right AUPRC |
|---|---|---|---|---|---|---|
| 5000 | 0.2 | 0.1 | 0.9680 ± 0.0428 | 0.9018 ± 0.1487 | 0.9740 ± 0.0209 | 0.9320 ± 0.0470 |
| 5000 | 0.2 | 0.3 | 0.9530 ± 0.0763 | 0.8864 ± 0.1433 | 0.9666 ± 0.0329 | 0.9099 ± 0.0701 |
| 5000 | 0.2 | 0.5 | 0.9544 ± 0.0443 | 0.8765 ± 0.1175 | 0.9653 ± 0.0486 | 0.9147 ± 0.1072 |
| 5000 | 0.3 | 0.1 | 0.9929 ± 0.0187 | 0.9818 ± 0.0463 | 0.9980 ± 0.0026 | 0.9940 ± 0.0053 |
| 5000 | 0.3 | 0.3 | 0.9944 ± 0.0080 | 0.9844 ± 0.0206 | 0.9964 ± 0.0059 | 0.9917 ± 0.0108 |
| 5000 | 0.3 | 0.5 | 0.9963 ± 0.0055 | 0.9915 ± 0.0090 | 0.9967 ± 0.0039 | 0.9916 ± 0.0079 |
| 5000 | 0.4 | 0.1 | 1.0000 ± 0.0001 | 0.9998 ± 0.0003 | 1.0000 ± 0.0001 | 0.9997 ± 0.0006 |
| 5000 | 0.4 | 0.3 | 0.9998 ± 0.0005 | 0.9996 ± 0.0010 | 1.0000 ± 0.0000 | 0.9997 ± 0.0003 |
| 5000 | 0.4 | 0.5 | 0.9995 ± 0.0014 | 0.9990 ± 0.0024 | 0.9996 ± 0.0016 | 0.9989 ± 0.0037 |
| 10000 | 0.2 | 0.1 | 0.9507 ± 0.0774 | 0.8060 ± 0.2014 | 0.9681 ± 0.0429 | 0.8566 ± 0.1472 |
| 10000 | 0.2 | 0.3 | 0.9565 ± 0.0666 | 0.8191 ± 0.2117 | 0.9592 ± 0.0432 | 0.8096 ± 0.1557 |
| 10000 | 0.2 | 0.5 | 0.9567 ± 0.0469 | 0.8145 ± 0.1607 | 0.9729 ± 0.0233 | 0.8574 ± 0.1144 |
| 10000 | 0.3 | 0.1 | 0.9972 ± 0.0064 | 0.9873 ± 0.0202 | 0.9764 ± 0.0900 | 0.9397 ± 0.2063 |
| 10000 | 0.3 | 0.3 | 0.9974 ± 0.0032 | 0.9867 ± 0.0128 | 0.9980 ± 0.0029 | 0.9882 ± 0.0114 |
| 10000 | 0.3 | 0.5 | 0.9958 ± 0.0069 | 0.9849 ± 0.0171 | 0.9948 ± 0.0147 | 0.9752 ± 0.0619 |
| 10000 | 0.4 | 0.1 | 1.0000 ± 0.0001 | 0.9995 ± 0.0008 | 1.0000 ± 0.0001 | 0.9995 ± 0.0006 |
| 10000 | 0.4 | 0.3 | 0.9998 ± 0.0005 | 0.9992 ± 0.0013 | 0.9996 ± 0.0012 | 0.9987 ± 0.0030 |
| 10000 | 0.4 | 0.5 | 0.9998 ± 0.0006 | 0.9990 ± 0.0022 | 0.9999 ± 0.0003 | 0.9995 ± 0.0007 |

### Supplementary Table S3. Performance of edgeR across all simulation settings.

| n_vars | p | r | Left AUROC | Left AUPRC | Right AUROC | Right AUPRC |
|---|---|---|---|---|---|---|
| 5000 | 0.2 | 0.1 | 0.9152 ± 0.1144 | 0.7109 ± 0.2912 | 0.8027 ± 0.1425 | 0.4538 ± 0.2600 |
| 5000 | 0.2 | 0.3 | 0.9714 ± 0.0387 | 0.8468 ± 0.1724 | 0.9423 ± 0.0799 | 0.7577 ± 0.2199 |
| 5000 | 0.2 | 0.5 | 0.9601 ± 0.0393 | 0.7324 ± 0.2159 | 0.9664 ± 0.0441 | 0.8202 ± 0.1795 |
| 5000 | 0.3 | 0.1 | 0.9938 ± 0.0131 | 0.9630 ± 0.0724 | 0.9522 ± 0.0536 | 0.7919 ± 0.1835 |
| 5000 | 0.3 | 0.3 | 0.9968 ± 0.0062 | 0.9747 ± 0.0462 | 0.9913 ± 0.0092 | 0.9398 ± 0.0597 |
| 5000 | 0.3 | 0.5 | 0.9939 ± 0.0077 | 0.9415 ± 0.0705 | 0.9987 ± 0.0030 | 0.9899 ± 0.0208 |
| 5000 | 0.4 | 0.1 | 0.9999 ± 0.0001 | 0.9995 ± 0.0010 | 0.9832 ± 0.0304 | 0.9147 ± 0.1291 |
| 5000 | 0.4 | 0.3 | 1.0000 ± 0.0001 | 0.9998 ± 0.0006 | 0.9994 ± 0.0011 | 0.9947 ± 0.0092 |
| 5000 | 0.4 | 0.5 | 0.9993 ± 0.0018 | 0.9926 ± 0.0185 | 1.0000 ± 0.0001 | 0.9998 ± 0.0005 |
| 10000 | 0.2 | 0.1 | 0.9011 ± 0.1266 | 0.5792 ± 0.2908 | 0.7474 ± 0.1754 | 0.2832 ± 0.2612 |
| 10000 | 0.2 | 0.3 | 0.9491 ± 0.1022 | 0.7409 ± 0.2726 | 0.8980 ± 0.1426 | 0.5823 ± 0.3028 |
| 10000 | 0.2 | 0.5 | 0.9632 ± 0.0368 | 0.6369 ± 0.2578 | 0.9754 ± 0.0510 | 0.8213 ± 0.2206 |
| 10000 | 0.3 | 0.1 | 0.9897 ± 0.0184 | 0.9040 ± 0.1330 | 0.9103 ± 0.1450 | 0.6626 ± 0.3192 |
| 10000 | 0.3 | 0.3 | 0.9976 ± 0.0053 | 0.9700 ± 0.0592 | 0.9969 ± 0.0037 | 0.9594 ± 0.0436 |
| 10000 | 0.3 | 0.5 | 0.9961 ± 0.0052 | 0.9279 ± 0.0901 | 0.9989 ± 0.0015 | 0.9815 ± 0.0252 |
| 10000 | 0.4 | 0.1 | 0.9997 ± 0.0006 | 0.9955 ± 0.0099 | 0.9921 ± 0.0168 | 0.9303 ± 0.1084 |
| 10000 | 0.4 | 0.3 | 1.0000 ± 0.0000 | 0.9998 ± 0.0005 | 0.9989 ± 0.0035 | 0.9864 ± 0.0411 |
| 10000 | 0.4 | 0.5 | 0.9993 ± 0.0022 | 0.9876 ± 0.0384 | 1.0000 ± 0.0000 | 0.9998 ± 0.0004 |

### Supplementary Table S4. Performance of DESeq2 across all simulation settings.

| n_vars | p | r | Left AUROC | Left AUPRC | Right AUROC | Right AUPRC |
|---|---|---|---|---|---|---|
| 5000 | 0.2 | 0.1 | 0.8882 ± 0.1331 | 0.6231 ± 0.2894 | 0.7516 ± 0.1531 | 0.3331 ± 0.1998 |
| 5000 | 0.2 | 0.3 | 0.9569 ± 0.0512 | 0.7744 ± 0.2041 | 0.9174 ± 0.0933 | 0.6472 ± 0.2173 |
| 5000 | 0.2 | 0.5 | 0.9377 ± 0.0572 | 0.6441 ± 0.2500 | 0.9459 ± 0.0580 | 0.7188 ± 0.1989 |
| 5000 | 0.3 | 0.1 | 0.9875 ± 0.0219 | 0.9149 ± 0.1140 | 0.9201 ± 0.0689 | 0.6192 ± 0.1990 |
| 5000 | 0.3 | 0.3 | 0.9926 ± 0.0109 | 0.9366 ± 0.0759 | 0.9804 ± 0.0162 | 0.8552 ± 0.0940 |
| 5000 | 0.3 | 0.5 | 0.9880 ± 0.0132 | 0.8919 ± 0.1022 | 0.9954 ± 0.0064 | 0.9565 ± 0.0452 |
| 5000 | 0.4 | 0.1 | 0.9989 ± 0.0008 | 0.9849 ± 0.0101 | 0.9626 ± 0.0450 | 0.7650 ± 0.1731 |
| 5000 | 0.4 | 0.3 | 0.9994 ± 0.0004 | 0.9901 ± 0.0073 | 0.9958 ± 0.0031 | 0.9510 ± 0.0261 |
| 5000 | 0.4 | 0.5 | 0.9981 ± 0.0032 | 0.9782 ± 0.0286 | 0.9989 ± 0.0005 | 0.9825 ± 0.0089 |
| 10000 | 0.2 | 0.1 | 0.8825 ± 0.1384 | 0.5143 ± 0.2847 | 0.7200 ± 0.1833 | 0.2388 ± 0.2295 |
| 10000 | 0.2 | 0.3 | 0.9386 ± 0.1139 | 0.6955 ± 0.2814 | 0.8834 ± 0.1518 | 0.5299 ± 0.2918 |
| 10000 | 0.2 | 0.5 | 0.9439 ± 0.0510 | 0.5369 ± 0.2616 | 0.9684 ± 0.0594 | 0.7732 ± 0.2262 |
| 10000 | 0.3 | 0.1 | 0.9850 ± 0.0247 | 0.8650 ± 0.1630 | 0.8964 ± 0.1552 | 0.6060 ± 0.3162 |
| 10000 | 0.3 | 0.3 | 0.9963 ± 0.0079 | 0.9552 ± 0.0817 | 0.9951 ± 0.0050 | 0.9328 ± 0.0560 |
| 10000 | 0.3 | 0.5 | 0.9923 ± 0.0095 | 0.8763 ± 0.1384 | 0.9980 ± 0.0025 | 0.9669 ± 0.0369 |
| 10000 | 0.4 | 0.1 | 0.9994 ± 0.0012 | 0.9889 ± 0.0173 | 0.9877 ± 0.0223 | 0.8778 ± 0.1441 |
| 10000 | 0.4 | 0.3 | 0.9999 ± 0.0001 | 0.9978 ± 0.0022 | 0.9983 ± 0.0048 | 0.9743 ± 0.0553 |
| 10000 | 0.4 | 0.5 | 0.9986 ± 0.0042 | 0.9757 ± 0.0631 | 0.9999 ± 0.0001 | 0.9969 ± 0.0044 |

### Supplementary Table S5. The 239 known AD risk genes collected from 33 GWAS studies in the NHGRI-EBI GWAS Catalog [8].

| Known AD risk gene | | | |
|---|---|---|---|
| *ABCA7* | *ABCA8* | *ABI3* | *AC006126.3* |
| *AC074212.3* | *ACE* | *ACSL6* | *ADAM10* |
| *ADAMTS1* | *ADAMTS20* | *ADAMTS4* | *AK128216* |
| *AKR7A3* | *ALPK2* | *ANKRD22* | *AOX1* |
| *AP2A2* | *APH1B* | *APOC1* | *APOE* |
| *ATXN7L1* | *BAALC* | *BANK1* | *BCAM* |
| *BCL3* | *BIN1* | *BZRAP1* | *BZRAP1-AS1* |
| *C17orf28* | *C1S* | *C1orf88* | *CACNA2D3* |
| *CADM2* | *CASS4* | *CBLC* | *CCZ1B* |
| *CD2AP* | *CD33* | *CDC42SE2* | *CDH1* |
| *CDON* | *CDR2L* | *CEACAM20* | *CEACAM22P* |
| *CELF1* | *CEP63* | *CETN4P* | *CHRNA2* |
| *CKAP5* | *CKM* | *CLASRP* | *CLDN18* |
| *CLEC3B* | *CLNK* | *CLPTM1* | *CLU* |
| *CNTNAP2* | *COBL* | *COL25A1* | *CPS1* |
| *CR1* | *CRHR1* | *CTB-171A8.1* | *CTB-179K24.3* |
| *CUGBP2* | *DAPL1* | *DISC1* | *DLC1* |
| *DNAH6* | *ECHDC3* | *EFL1* | *EGLN3* |
| *ELL* | *EPHA1* | *EXOC3L2* | *F2* |
| *FAM113B* | *FAM155A* | *FAM172BP* | *FAM19A5* |
| *FAM20C* | *FAM47E* | *FAT3* | *FBXL7* |
| *FBXO40* | *FERMT2* | *FLJ37543* | *FNIP1* |
| *FST* | *GAPDHP15* | *GEMIN7* | *GLTSCR1* |
| *GPR180* | *GTF2H3* | *HBEGF* | *HDAC9* |
| *HESX1* | *HLA-DRA* | *HLA-DRB1* | *HLA-DRB5* |
| *HMCN1* | *HMGA2* | *HS3ST1* | *IGH* |
| *IGSF23* | *IL21* | *IMMP2L* | *INPP5D* |
| *INPPD5* | *IQCK* | *IQGAP2* | *IRAK1BP1* |
| *IRF6* | *ISYNA1* | *JPH3* | *KANSL1* |
| *KAT8* | *KCNN3* | *KCTD8* | *KRT18P45* |
| *KSR2* | *LINC00290* | *LINC00899* | *LOC257396* |
| *LRRC27A* | *LUZP2* | *MACROD2* | *MAF* |
| *MAPT* | *MARK4* | *MECOM* | *MEF2C* |
| *MGC45800* | *MGMT* | *MIR142* | *MIR5582* |
| *MKI67* | *MMP12* | *MMP3* | *MPZL1* |
| *MS4* | *MS4A4A* | *MS4A6A* | *MS4A6E* |
| *MSRA* | *MTSS1L* | *NDUFAF6* | *NEGR1* |
| *NEGR1-IT1* | *NEK10* | *NFIC* | *NHLRC3* |
| *NKPD1* | *NME8* | *NPVF* | *NTM* |
| *OARD1* | *ODZ4* | *OFCC1* | *OR2B2* |
| *OR7E89P* | *PCDH7* | *PCNX* | *PDE1A* |
| *PEX6* | *PFDN1* | *PGBD3P4* | *PICALM* |
| *PMS2CL* | *POLN* | *PPIAP17* | *PPP1R3B* |
| *PROSER1* | *PTK2B* | *PVR* | *PVRL2* |
| *RAPGEF6* | *RASSF5* | *RBBP4P4* | *RELB* |
| *RELN* | *RIN3* | *RPL13AP7* | *RPL3P8* |
| *RPL7AP36* | *RPS27P10* | *SCAPER* | *SCIMP* |
| *SEC24B* | *SELO* | *SELP* | *SENP7* |
| *SERINC5* | *SESTD1* | *SH2D4B* | *SHANK2* |
| *SLC10A2* | *SLC24A4* | *SLC4A1AP* | *SLMAP* |
| *SORD* | *SORL1* | *SOX14* | *SPPL2A* |
| *SQSTM1* | *SSBP4* | *STAG3* | *STAU2* |
| *STK24* | *SUCLG2P4* | *SUZ12P1* | *SYNGAP1* |
| *TAS2R5* | *TAS2R62P* | *TBXAS1* | *TFEB* |
| *THSD7A* | *TIMP2* | *TMCO4* | *TMEM106B* |
| *TOMM40* | *TREM2* | *TREML2* | *TRIP4* |
| *TRPM1* | *TSPAN14* | *TSPOAP1-AS1* | *UBA52P1* |
| *UBE2D2* | *UGT1A10* | *UGT1A8* | *USP6NL* |
| *VN2R20P* | *WNT3* | *WWOX* | *ZCWPW1* |
| *ZNF12* | *ZNF224* | *snoZ6* | |

### Supplementary Table S6. Sizes and compositions of ADBrainNexus network modules.

The Louvain algorithm implemented in NetworkX [9] was used to partition the ADBrainNexus [10] subnetwork constructed from the top 200 TailEnrich-specific genes and the 239 known AD risk genes by retaining edges with weights greater than 0.75. Four modules containing at least 15 of the top 200 TailEnrich-specific genes were retained for enrichment analysis. The table reports the sizes of these four modules (M1, M2, M3, and M4) and the genes in each module.

| Module | Module size | Genes |
|---|---|---|
| M1 | 36 | *AP2A2, ASAP3, CBLC, CD2AP, DISC1, HBEGF, IGF1R, IMPA2, INPP5D, ISYNA1, ITSN1, MAF, MYO10, PDE1A, PDE4B, PIK3C2A, PLCD3, PLCG1, PPP1R3B, PTK2B, RAB27A, RAB31, RAPGEF6, RASSF5, RELN, RHOQ, SEC14L1, SERINC5, SH3BP2, SORBS1, STAG3, TCF12, THSD7A, TIMP2, UBE2D2, UTRN* |
| M2 | 32 | *ADAM10, APC2, ARID1A, BCAM, CDKN2C, CHRNA2, COBL, CRHR1, ENAH, EPHA1, F2, FERMT2, FOXN3, FZD1, HMGA2, IQGAP2, JPH3, KAT6A, LMAN1, LPIN1, NEBL, PEAK1, PON2, ROCK1, SEMA4B, SLC4A1AP, SNAP23, SOX2, TMEM106B, USP6NL, WNT3, YES1* |
| M3 | 28 | *ABCA7, ANKFY1, APOC1, APOE, BCL3, BIN1, BNIP2, CLU, DLC1, DMPK, EGLN3, EMP2, FLCN, MAP3K5, OTUD6B, PICALM, PITPNC1, PLEKHF2, PNPLA8, SCAMP2, SEC24B, SH3GLB1, SLC3A2, STK24, TMED10, TMEM123, TMEM170A, TNFRSF10B* |
| M4 | 24 | *ABCA8, ABCC9, ACSL6, ATP6V0E1, CALCOCO2, CEP350, CLEC3B, HSPA5, IDH1, IMMP2L, MSI2, NDUFAF6, PEX2, PEX6, POLN, SELENBP1, SLC30A1, SLC41A1, SLC7A1, SQSTM1, TCF3, TOMM40, VPS13C, ZMIZ1* |

### Supplementary Table S7. The expression-trait correlation results on the MSBB-BM36 dataset.

The table reports correlation results for the 225 genes that met the criteria for at least one of the three AD-related traits. Pearson r, p values and FDR are shown for Braak stage, CDR, and CERAD scores.

| Gene | Trait | r | p | FDR | Gene | Trait | r | p | FDR |
|---|---|---|---|---|---|---|---|---|---|
| PHYHD1 | Braak | 0.402 | 1.52e-08 | 1.84e-06 | PHYHD1 | CDR | 0.317 | 5.08e-06 | 2.94e-05 |
| PHYHD1 | CERAD | 0.406 | 3.62e-07 | 7.79e-06 | NEBL | Braak | 0.267 | 2.39e-04 | 8.43e-04 |
| NEBL | CDR | 0.235 | 8.17e-04 | 1.87e-03 | NEBL | CERAD | 0.379 | 2.32e-06 | 2.64e-05 |
| OGFRL1 | Braak | 0.259 | 3.79e-04 | 1.20e-03 | OGFRL1 | CDR | 0.161 | 0.023 | 3.76e-02 |
| OGFRL1 | CERAD | 0.377 | 2.56e-06 | 2.84e-05 | MRAS | Braak | 0.335 | 3.18e-06 | 3.94e-05 |
| MRAS | CDR | 0.260 | 2.05e-04 | 5.82e-04 | MRAS | CERAD | 0.287 | 4.30e-04 | 1.32e-03 |
| PLCD3 | Braak | 0.309 | 1.94e-05 | 1.28e-04 | PLCD3 | CDR | 0.289 | 3.47e-05 | 1.43e-04 |
| PLCD3 | CERAD | 0.367 | 5.11e-06 | 4.70e-05 | PAX6 | Braak | 0.249 | 6.31e-04 | 1.85e-03 |
| PAX6 | CDR | 0.326 | 2.58e-06 | 1.80e-05 | PAX6 | CERAD | 0.318 | 8.70e-05 | 3.71e-04 |
| EPHX1 | Braak | 0.278 | 1.33e-04 | 5.38e-04 | EPHX1 | CDR | 0.329 | 2.08e-06 | 1.53e-05 |
| EPHX1 | CERAD | 0.347 | 1.70e-05 | 1.09e-04 | RHOQ | Braak | 0.351 | 1.02e-06 | 1.53e-05 |
| RHOQ | CDR | 0.270 | 1.16e-04 | 3.78e-04 | RHOQ | CERAD | 0.396 | 7.15e-07 | 1.24e-05 |
| SOX9 | Braak | 0.382 | 8.63e-08 | 3.39e-06 | SOX9 | CDR | 0.351 | 3.76e-07 | 4.04e-06 |
| SOX9 | CERAD | 0.398 | 6.24e-07 | 1.14e-05 | CLU | Braak | 0.206 | 0.005 | 1.05e-02 |
| CLU | CDR | 0.281 | 5.85e-05 | 2.18e-04 | CLU | CERAD | 0.317 | 9.21e-05 | 3.85e-04 |
| EMP2 | Braak | 0.322 | 7.77e-06 | 6.34e-05 | EMP2 | CDR | 0.396 | 7.59e-09 | 2.57e-07 |
| EMP2 | CERAD | 0.340 | 2.68e-05 | 1.54e-04 | DLC1 | Braak | 0.359 | 5.28e-07 | 1.09e-05 |
| DLC1 | CDR | 0.257 | 2.53e-04 | 6.87e-04 | DLC1 | CERAD | 0.275 | 7.76e-04 | 2.11e-03 |
| RNF19A | Braak | 0.130 | 0.076 | 1.07e-01 | RNF19A | CDR | 0.158 | 0.026 | 4.09e-02 |
| RNF19A | CERAD | 0.307 | 1.58e-04 | 5.73e-04 | YES1 | Braak | 0.343 | 1.77e-06 | 2.48e-05 |
| YES1 | CDR | 0.323 | 3.40e-06 | 2.21e-05 | YES1 | CERAD | 0.309 | 1.47e-04 | 5.42e-04 |
| SORBS1 | Braak | 0.367 | 2.93e-07 | 6.75e-06 | SORBS1 | CDR | 0.402 | 4.32e-09 | 1.99e-07 |
| SORBS1 | CERAD | 0.473 | 1.54e-09 | 4.98e-07 | PLEKHO2 | Braak | 0.332 | 4.06e-06 | 4.44e-05 |
| PLEKHO2 | CDR | 0.400 | 5.18e-09 | 2.22e-07 | PLEKHO2 | CERAD | 0.328 | 5.26e-05 | 2.56e-04 |
| CHDH | Braak | 0.379 | 1.08e-07 | 3.39e-06 | CHDH | CDR | 0.332 | 1.72e-06 | 1.31e-05 |
| CHDH | CERAD | 0.370 | 4.05e-06 | 4.15e-05 | GNA12 | Braak | 0.409 | 7.76e-09 | 1.25e-06 |
| GNA12 | CDR | 0.417 | 9.55e-10 | 7.70e-08 | GNA12 | CERAD | 0.437 | 3.35e-08 | 2.70e-06 |
| ENAH | Braak | 0.306 | 2.28e-05 | 1.42e-04 | ENAH | CDR | 0.311 | 8.15e-06 | 4.34e-05 |
| ENAH | CERAD | 0.414 | 1.94e-07 | 5.74e-06 | ARL6IP6 | Braak | 0.339 | 2.50e-06 | 3.22e-05 |
| ARL6IP6 | CDR | 0.253 | 3.14e-04 | 8.23e-04 | ARL6IP6 | CERAD | 0.326 | 5.90e-05 | 2.77e-04 |
| TMEM170A | Braak | 0.318 | 1.05e-05 | 7.94e-05 | TMEM170A | CDR | 0.317 | 5.04e-06 | 2.94e-05 |
| TMEM170A | CERAD | 0.313 | 1.18e-04 | 4.61e-04 | ECE1 | Braak | 0.330 | 4.46e-06 | 4.76e-05 |
| ECE1 | CDR | 0.326 | 2.61e-06 | 1.80e-05 | ECE1 | CERAD | 0.389 | 1.21e-06 | 1.66e-05 |
| CDH20 | Braak | 0.305 | 2.51e-05 | 1.53e-04 | CDH20 | CDR | 0.319 | 4.61e-06 | 2.75e-05 |
| CDH20 | CERAD | 0.305 | 1.80e-04 | 6.33e-04 | WSCD1 | Braak | 0.289 | 6.85e-05 | 3.20e-04 |
| WSCD1 | CDR | 0.344 | 6.82e-07 | 6.76e-06 | WSCD1 | CERAD | 0.251 | 0.002 | 4.94e-03 |
| SLC41A1 | Braak | 0.293 | 5.32e-05 | 2.60e-04 | SLC41A1 | CDR | 0.311 | 7.66e-06 | 4.11e-05 |
| SLC41A1 | CERAD | 0.334 | 3.63e-05 | 1.93e-04 | MAPKAPK2 | Braak | 0.342 | 2.00e-06 | 2.73e-05 |
| MAPKAPK2 | CDR | 0.322 | 3.57e-06 | 2.30e-05 | MAPKAPK2 | CERAD | 0.348 | 1.66e-05 | 1.08e-04 |
| MSI2 | Braak | 0.397 | 2.24e-08 | 1.84e-06 | MSI2 | CDR | 0.422 | 5.58e-10 | 5.99e-08 |
| MSI2 | CERAD | 0.418 | 1.48e-07 | 5.74e-06 | TRIP6 | Braak | 0.323 | 7.59e-06 | 6.27e-05 |
| TRIP6 | CDR | 0.386 | 1.82e-08 | 5.09e-07 | TRIP6 | CERAD | 0.402 | 4.70e-07 | 9.47e-06 |
| KLF3 | Braak | 0.281 | 1.09e-04 | 4.66e-04 | KLF3 | CDR | 0.294 | 2.51e-05 | 1.10e-04 |
| KLF3 | CERAD | 0.336 | 3.30e-05 | 1.80e-04 | MFAP3L | Braak | 0.192 | 0.009 | 1.67e-02 |
| MFAP3L | CDR | 0.253 | 3.08e-04 | 8.12e-04 | MFAP3L | CERAD | 0.352 | 1.30e-05 | 8.75e-05 |
| TBL1X | Braak | 0.449 | 1.63e-10 | 1.05e-07 | TBL1X | CDR | 0.396 | 7.49e-09 | 2.58e-07 |
| TBL1X | CERAD | 0.416 | 1.67e-07 | 5.74e-06 | KMT2C | Braak | 0.376 | 1.37e-07 | 3.84e-06 |
| KMT2C | CDR | 0.404 | 3.29e-09 | 1.63e-07 | KMT2C | CERAD | 0.451 | 1.09e-08 | 1.75e-06 |
| TCF12 | Braak | 0.297 | 4.17e-05 | 2.18e-04 | TCF12 | CDR | 0.298 | 1.94e-05 | 8.91e-05 |
| TCF12 | CERAD | 0.356 | 9.78e-06 | 7.06e-05 | ARID1A | Braak | 0.302 | 2.92e-05 | 1.69e-04 |
| ARID1A | CDR | 0.281 | 5.96e-05 | 2.18e-04 | ARID1A | CERAD | 0.319 | 8.66e-05 | 3.71e-04 |
| ANKFY1 | Braak | 0.336 | 2.92e-06 | 3.69e-05 | ANKFY1 | CDR | 0.370 | 7.46e-08 | 1.17e-06 |
| ANKFY1 | CERAD | 0.434 | 4.35e-08 | 3.12e-06 | NEO1 | Braak | 0.355 | 7.30e-07 | 1.30e-05 |
| NEO1 | CDR | 0.285 | 4.58e-05 | 1.82e-04 | NEO1 | CERAD | 0.382 | 1.86e-06 | 2.30e-05 |
| MSRB3 | Braak | 0.344 | 1.66e-06 | 2.43e-05 | MSRB3 | CDR | 0.345 | 6.39e-07 | 6.43e-06 |
| MSRB3 | CERAD | 0.360 | 7.97e-06 | 6.12e-05 | MTMR3 | Braak | 0.380 | 1.03e-07 | 3.39e-06 |
| MTMR3 | CDR | 0.398 | 6.07e-09 | 2.40e-07 | MTMR3 | CERAD | 0.422 | 1.06e-07 | 5.26e-06 |
| CD151 | Braak | 0.278 | 1.27e-04 | 5.20e-04 | CD151 | CDR | 0.350 | 4.18e-07 | 4.41e-06 |
| CD151 | CERAD | 0.349 | 1.56e-05 | 1.02e-04 | ROCK1 | Braak | 0.299 | 3.62e-05 | 1.99e-04 |
| ROCK1 | CDR | 0.347 | 5.11e-07 | 5.31e-06 | ROCK1 | CERAD | 0.354 | 1.14e-05 | 8.01e-05 |
| ERLIN2 | Braak | 0.186 | 0.011 | 2.10e-02 | ERLIN2 | CDR | 0.249 | 3.93e-04 | 9.88e-04 |
| ERLIN2 | CERAD | 0.309 | 1.41e-04 | 5.28e-04 | LPIN1 | Braak | 0.371 | 2.06e-07 | 5.10e-06 |
| LPIN1 | CDR | 0.360 | 1.78e-07 | 2.24e-06 | LPIN1 | CERAD | 0.399 | 5.94e-07 | 1.12e-05 |
| SEMA4B | Braak | 0.309 | 1.93e-05 | 1.28e-04 | SEMA4B | CDR | 0.274 | 9.14e-05 | 3.11e-04 |
| SEMA4B | CERAD | 0.284 | 4.94e-04 | 1.48e-03 | BNIP2 | Braak | 0.270 | 2.07e-04 | 7.59e-04 |
| BNIP2 | CDR | 0.282 | 5.41e-05 | 2.05e-04 | BNIP2 | CERAD | 0.349 | 1.51e-05 | 1.00e-04 |
| NFIA | Braak | 0.380 | 1.03e-07 | 3.39e-06 | NFIA | CDR | 0.340 | 9.38e-07 | 8.20e-06 |
| NFIA | CERAD | 0.379 | 2.34e-06 | 2.64e-05 | CARD8 | Braak | 0.352 | 9.54e-07 | 1.48e-05 |

| Gene | Trait | r | p | FDR | Gene | Trait | r | p | FDR |
|---|---|---|---|---|---|---|---|---|---|
| CARD8 | CDR | 0.368 | 9.43e-08 | 1.38e-06 | CARD8 | CERAD | 0.380 | 2.13e-06 | 2.55e-05 |
| SCAMP2 | Braak | 0.305 | 2.52e-05 | 1.53e-04 | SCAMP2 | CDR | 0.378 | 3.78e-08 | 7.17e-07 |
| SCAMP2 | CERAD | 0.369 | 4.55e-06 | 4.38e-05 | ZHX3 | Braak | 0.330 | 4.50e-06 | 4.76e-05 |
| ZHX3 | CDR | 0.328 | 2.29e-06 | 1.62e-05 | ZHX3 | CERAD | 0.333 | 3.94e-05 | 2.03e-04 |
| BMPR1A | Braak | 0.215 | 0.003 | 7.64e-03 | BMPR1A | CDR | 0.183 | 0.010 | 1.69e-02 |
| BMPR1A | CERAD | 0.307 | 1.61e-04 | 5.79e-04 | FYN | Braak | 0.333 | 3.61e-06 | 4.15e-05 |
| FYN | CDR | 0.376 | 4.59e-08 | 8.37e-07 | FYN | CERAD | 0.413 | 2.14e-07 | 5.74e-06 |
| TMED10 | Braak | 0.205 | 0.005 | 1.07e-02 | TMED10 | CDR | 0.231 | 0.001 | 2.29e-03 |
| TMED10 | CERAD | 0.306 | 1.71e-04 | 6.11e-04 | MLC1 | Braak | 0.275 | 1.58e-04 | 6.12e-04 |
| MLC1 | CDR | 0.310 | 8.23e-06 | 4.35e-05 | MLC1 | CERAD | 0.281 | 5.81e-04 | 1.67e-03 |
| COLGALT1 | Braak | 0.286 | 7.89e-05 | 3.55e-04 | COLGALT1 | CDR | 0.307 | 1.03e-05 | 5.18e-05 |
| COLGALT1 | CERAD | 0.276 | 7.38e-04 | 2.02e-03 | ASAP3 | Braak | 0.296 | 4.40e-05 | 2.21e-04 |
| ASAP3 | CDR | 0.244 | 5.11e-04 | 1.23e-03 | ASAP3 | CERAD | 0.332 | 4.16e-05 | 2.09e-04 |
| BMP7 | Braak | 0.270 | 2.05e-04 | 7.57e-04 | BMP7 | CDR | 0.303 | 1.37e-05 | 6.78e-05 |
| BMP7 | CERAD | 0.270 | 9.48e-04 | 2.48e-03 | TNFRSF10B | Braak | 0.270 | 2.05e-04 | 7.57e-04 |
| TNFRSF10B | CDR | 0.339 | 9.67e-07 | 8.20e-06 | TNFRSF10B | CERAD | 0.248 | 0.002 | 5.35e-03 |
| CDC42SE1 | Braak | 0.324 | 7.15e-06 | 6.26e-05 | CDC42SE1 | CDR | 0.318 | 4.69e-06 | 2.77e-05 |
| CDC42SE1 | CERAD | 0.310 | 1.36e-04 | 5.12e-04 | UTRN | Braak | 0.358 | 5.73e-07 | 1.15e-05 |
| UTRN | CDR | 0.339 | 9.59e-07 | 8.20e-06 | UTRN | CERAD | 0.420 | 1.24e-07 | 5.63e-06 |
| WBP1L | Braak | 0.323 | 7.44e-06 | 6.27e-05 | WBP1L | CDR | 0.363 | 1.39e-07 | 1.90e-06 |
| WBP1L | CERAD | 0.391 | 1.00e-06 | 1.46e-05 | CSRP1 | Braak | 0.296 | 4.28e-05 | 2.19e-04 |
| CSRP1 | CDR | 0.302 | 1.52e-05 | 7.25e-05 | CSRP1 | CERAD | 0.341 | 2.52e-05 | 1.49e-04 |
| TCF3 | Braak | 0.354 | 8.03e-07 | 1.34e-05 | TCF3 | CDR | 0.319 | 4.39e-06 | 2.73e-05 |
| TCF3 | CERAD | 0.290 | 3.78e-04 | 1.19e-03 | IKBKB | Braak | 0.400 | 1.81e-08 | 1.84e-06 |
| IKBKB | CDR | 0.398 | 6.34e-09 | 2.40e-07 | IKBKB | CERAD | 0.396 | 7.31e-07 | 1.24e-05 |
| SH3BP2 | Braak | 0.381 | 8.97e-08 | 3.39e-06 | SH3BP2 | CDR | 0.293 | 2.69e-05 | 1.16e-04 |
| SH3BP2 | CERAD | 0.381 | 2.06e-06 | 2.51e-05 | SNAP23 | Braak | 0.298 | 3.95e-05 | 2.08e-04 |
| SNAP23 | CDR | 0.288 | 3.74e-05 | 1.52e-04 | SNAP23 | CERAD | 0.325 | 6.15e-05 | 2.83e-04 |
| SEC14L2 | Braak | 0.274 | 1.62e-04 | 6.27e-04 | SEC14L2 | CDR | 0.332 | 1.66e-06 | 1.28e-05 |
| SEC14L2 | CERAD | 0.342 | 2.31e-05 | 1.38e-04 | KAT6A | Braak | 0.383 | 7.84e-08 | 3.39e-06 |
| KAT6A | CDR | 0.310 | 8.57e-06 | 4.45e-05 | KAT6A | CERAD | 0.441 | 2.41e-08 | 2.58e-06 |
| ZMIZ1 | Braak | 0.174 | 0.018 | 3.12e-02 | ZMIZ1 | CDR | 0.290 | 3.28e-05 | 1.38e-04 |
| ZMIZ1 | CERAD | 0.326 | 5.62e-05 | 2.72e-04 | WDFY2 | Braak | 0.393 | 3.28e-08 | 2.12e-06 |
| WDFY2 | CDR | 0.407 | 2.54e-09 | 1.36e-07 | WDFY2 | CERAD | 0.475 | 1.30e-09 | 4.98e-07 |
| ZC3H11A | Braak | 0.307 | 2.13e-05 | 1.34e-04 | ZC3H11A | CDR | 0.227 | 0.001 | 2.79e-03 |
| ZC3H11A | CERAD | 0.313 | 1.18e-04 | 4.61e-04 | PEAK1 | Braak | 0.303 | 2.85e-05 | 1.68e-04 |
| PEAK1 | CDR | 0.268 | 1.30e-04 | 4.12e-04 | PEAK1 | CERAD | 0.356 | 9.86e-06 | 7.06e-05 |
| MOB3C | Braak | 0.365 | 3.33e-07 | 7.40e-06 | MOB3C | CDR | 0.375 | 5.10e-08 | 8.75e-07 |
| MOB3C | CERAD | 0.344 | 2.05e-05 | 1.25e-04 | MAT2A | Braak | 0.315 | 1.25e-05 | 9.27e-05 |
| MAT2A | CDR | 0.247 | 4.37e-04 | 1.08e-03 | MAT2A | CERAD | 0.339 | 2.76e-05 | 1.57e-04 |
| LMO2 | Braak | 0.233 | 0.001 | 3.57e-03 | LMO2 | CDR | 0.319 | 4.55e-06 | 2.75e-05 |
| LMO2 | CERAD | 0.322 | 7.31e-05 | 3.23e-04 | FGD4 | Braak | 0.217 | 0.003 | 6.92e-03 |
| FGD4 | CDR | 0.241 | 6.11e-04 | 1.43e-03 | FGD4 | CERAD | 0.304 | 1.85e-04 | 6.47e-04 |
| DMPK | Braak | 0.325 | 6.42e-06 | 5.87e-05 | DMPK | CDR | 0.341 | 8.26e-07 | 7.72e-06 |
| DMPK | CERAD | 0.231 | 0.005 | 9.41e-03 | RAB27A | Braak | 0.260 | 3.65e-04 | 1.17e-03 |
| RAB27A | CDR | 0.181 | 0.011 | 1.85e-02 | RAB27A | CERAD | 0.337 | 3.13e-05 | 1.73e-04 |
| CALCOCO2 | Braak | 0.308 | 2.00e-05 | 1.31e-04 | CALCOCO2 | CDR | 0.312 | 7.25e-06 | 3.96e-05 |
| CALCOCO2 | CERAD | 0.371 | 3.93e-06 | 4.15e-05 | PRPF38B | Braak | 0.251 | 5.62e-04 | 1.70e-03 |
| PRPF38B | CDR | 0.240 | 6.59e-04 | 1.54e-03 | PRPF38B | CERAD | 0.334 | 3.79e-05 | 1.98e-04 |
| NHSL1 | Braak | 0.394 | 3.06e-08 | 2.11e-06 | NHSL1 | CDR | 0.395 | 8.06e-09 | 2.60e-07 |
| NHSL1 | CERAD | 0.391 | 1.03e-06 | 1.48e-05 | MAOB | Braak | 0.246 | 7.39e-04 | 2.12e-03 |
| MAOB | CDR | 0.250 | 3.62e-04 | 9.20e-04 | MAOB | CERAD | 0.400 | 5.40e-07 | 1.05e-05 |
| SFT2D2 | Braak | 0.287 | 7.66e-05 | 3.50e-04 | SFT2D2 | CDR | 0.347 | 5.43e-07 | 5.55e-06 |
| SFT2D2 | CERAD | 0.383 | 1.76e-06 | 2.22e-05 | HIPK1 | Braak | 0.320 | 9.24e-06 | 7.18e-05 |
| HIPK1 | CDR | 0.293 | 2.75e-05 | 1.18e-04 | HIPK1 | CERAD | 0.410 | 2.57e-07 | 6.28e-06 |
| NES | Braak | 0.382 | 8.15e-08 | 3.39e-06 | NES | CDR | 0.430 | 2.40e-10 | 3.87e-08 |
| NES | CERAD | 0.417 | 1.63e-07 | 5.74e-06 | STK38 | Braak | 0.334 | 3.54e-06 | 4.15e-05 |
| STK38 | CDR | 0.409 | 2.18e-09 | 1.27e-07 | STK38 | CERAD | 0.325 | 6.11e-05 | 2.83e-04 |
| IFI35 | Braak | 0.269 | 2.19e-04 | 7.94e-04 | IFI35 | CDR | 0.303 | 1.40e-05 | 6.81e-05 |
| IFI35 | CERAD | 0.286 | 4.46e-04 | 1.36e-03 | ARHGEF10L | Braak | 0.314 | 1.36e-05 | 9.98e-05 |
| ARHGEF10L | CDR | 0.300 | 1.73e-05 | 8.04e-05 | ARHGEF10L | CERAD | 0.316 | 9.94e-05 | 4.11e-04 |
| ZNF532 | Braak | 0.368 | 2.69e-07 | 6.42e-06 | ZNF532 | CDR | 0.342 | 7.85e-07 | 7.44e-06 |
| ZNF532 | CERAD | 0.388 | 1.28e-06 | 1.71e-05 | PLD2 | Braak | 0.270 | 2.01e-04 | 7.52e-04 |
| PLD2 | CDR | 0.268 | 1.28e-04 | 4.07e-04 | PLD2 | CERAD | 0.309 | 1.46e-04 | 5.40e-04 |
| USP47 | Braak | 0.203 | 0.006 | 1.16e-02 | USP47 | CDR | 0.141 | 0.047 | 6.94e-02 |
| USP47 | CERAD | 0.322 | 7.04e-05 | 3.17e-04 | REST | Braak | 0.328 | 5.32e-06 | 5.19e-05 |
| REST | CDR | 0.351 | 3.77e-07 | 4.04e-06 | REST | CERAD | 0.376 | 2.80e-06 | 3.05e-05 |
| NINJ1 | Braak | 0.210 | 0.004 | 9.04e-03 | NINJ1 | CDR | 0.283 | 5.34e-05 | 2.03e-04 |
| NINJ1 | CERAD | 0.319 | 8.56e-05 | 3.71e-04 | DOK1 | Braak | 0.241 | 9.81e-04 | 2.66e-03 |
| DOK1 | CDR | 0.361 | 1.68e-07 | 2.16e-06 | DOK1 | CERAD | 0.264 | 0.001 | 3.10e-03 |
| BCL2 | Braak | 0.374 | 1.66e-07 | 4.46e-06 | BCL2 | CDR | 0.382 | 2.72e-08 | 6.27e-07 |
| BCL2 | CERAD | 0.406 | 3.44e-07 | 7.64e-06 | DPYSL3 | Braak | 0.323 | 7.52e-06 | 6.27e-05 |
| DPYSL3 | CDR | 0.340 | 9.31e-07 | 8.20e-06 | DPYSL3 | CERAD | 0.413 | 2.13e-07 | 5.74e-06 |
| ZSWIM6 | Braak | 0.220 | 0.003 | 6.15e-03 | ZSWIM6 | CDR | 0.212 | 0.003 | 5.24e-03 |
| ZSWIM6 | CERAD | 0.313 | 1.17e-04 | 4.61e-04 | FAH | Braak | 0.212 | 0.004 | 8.42e-03 |
| FAH | CDR | 0.309 | 9.29e-06 | 4.75e-05 | FAH | CERAD | 0.282 | 5.53e-04 | 1.61e-03 |
| TCIRG1 | Braak | 0.300 | 3.38e-05 | 1.91e-04 | TCIRG1 | CDR | 0.342 | 7.58e-07 | 7.29e-06 |
| TCIRG1 | CERAD | 0.269 | 9.99e-04 | 2.59e-03 | TPCN1 | Braak | 0.325 | 6.39e-06 | 5.87e-05 |

| Gene | Trait | r | p | FDR | Gene | Trait | r | p | FDR |
|---|---|---|---|---|---|---|---|---|---|
| TPCN1 | CDR | 0.281 | 6.00e-05 | 2.18e-04 | TPCN1 | CERAD | 0.314 | 1.09e-04 | 4.43e-04 |
| NXN | Braak | 0.237 | 0.001 | 3.07e-03 | NXN | CDR | 0.367 | 1.00e-07 | 1.43e-06 |
| NXN | CERAD | 0.216 | 0.009 | 1.53e-02 | GALNT10 | Braak | 0.324 | 7.19e-06 | 6.26e-05 |
| GALNT10 | CDR | 0.383 | 2.39e-08 | 6.20e-07 | GALNT10 | CERAD | 0.392 | 9.77e-07 | 1.46e-05 |
| MED13L | Braak | 0.268 | 2.29e-04 | 8.23e-04 | MED13L | CDR | 0.249 | 3.94e-04 | 9.88e-04 |
| MED13L | CERAD | 0.324 | 6.30e-05 | 2.88e-04 | ACVR1 | Braak | 0.243 | 8.63e-04 | 2.38e-03 |
| ACVR1 | CDR | 0.301 | 1.59e-05 | 7.51e-05 | ACVR1 | CERAD | 0.389 | 1.19e-06 | 1.66e-05 |
| DCAF12 | Braak | 0.257 | 4.12e-04 | 1.29e-03 | DCAF12 | CDR | 0.265 | 1.56e-04 | 4.73e-04 |
| DCAF12 | CERAD | 0.368 | 4.63e-06 | 4.38e-05 | TBC1D2B | Braak | 0.383 | 7.72e-08 | 3.39e-06 |
| TBC1D2B | CDR | 0.432 | 1.89e-10 | 3.87e-08 | TBC1D2B | CERAD | 0.411 | 2.52e-07 | 6.29e-06 |
| PELI2 | Braak | 0.379 | 1.10e-07 | 3.39e-06 | PELI2 | CDR | 0.289 | 3.43e-05 | 1.42e-04 |
| PELI2 | CERAD | 0.392 | 9.76e-07 | 1.46e-05 | SOWAHC | Braak | 0.305 | 2.42e-05 | 1.50e-04 |
| SOWAHC | CDR | 0.268 | 1.29e-04 | 4.11e-04 | SOWAHC | CERAD | 0.299 | 2.41e-04 | 8.17e-04 |
| TMEM43 | Braak | 0.292 | 5.46e-05 | 2.61e-04 | TMEM43 | CDR | 0.383 | 2.45e-08 | 6.20e-07 |
| TMEM43 | CERAD | 0.345 | 1.93e-05 | 1.20e-04 | TULP3 | Braak | 0.299 | 3.67e-05 | 2.00e-04 |
| TULP3 | CDR | 0.294 | 2.52e-05 | 1.10e-04 | TULP3 | CERAD | 0.313 | 1.17e-04 | 4.61e-04 |
| APPL2 | Braak | 0.317 | 1.09e-05 | 8.18e-05 | APPL2 | CDR | 0.379 | 3.65e-08 | 7.12e-07 |
| APPL2 | CERAD | 0.457 | 6.56e-09 | 1.40e-06 | VANGL2 | Braak | 0.354 | 8.13e-07 | 1.34e-05 |
| VANGL2 | CDR | 0.319 | 4.46e-06 | 2.73e-05 | VANGL2 | CERAD | 0.386 | 1.43e-06 | 1.88e-05 |
| DHRS3 | Braak | 0.183 | 0.013 | 2.34e-02 | DHRS3 | CDR | 0.368 | 9.44e-08 | 1.38e-06 |
| DHRS3 | CERAD | 0.280 | 6.00e-04 | 1.69e-03 | APOL3 | Braak | 0.325 | 6.46e-06 | 5.87e-05 |
| APOL3 | CDR | 0.370 | 7.89e-08 | 1.21e-06 | APOL3 | CERAD | 0.363 | 6.50e-06 | 5.30e-05 |
| AHCYL1 | Braak | 0.184 | 0.012 | 2.25e-02 | AHCYL1 | CDR | 0.254 | 2.92e-04 | 7.78e-04 |
| AHCYL1 | CERAD | 0.333 | 4.00e-05 | 2.04e-04 | AXL | Braak | 0.333 | 3.81e-06 | 4.23e-05 |
| AXL | CDR | 0.301 | 1.62e-05 | 7.59e-05 | AXL | CERAD | 0.340 | 2.57e-05 | 1.50e-04 |
| VEZF1 | Braak | 0.299 | 3.61e-05 | 1.99e-04 | VEZF1 | CDR | 0.267 | 1.36e-04 | 4.24e-04 |
| VEZF1 | CERAD | 0.359 | 8.43e-06 | 6.25e-05 | RAB13 | Braak | 0.273 | 1.69e-04 | 6.46e-04 |
| RAB13 | CDR | 0.343 | 7.06e-07 | 6.89e-06 | RAB13 | CERAD | 0.364 | 6.08e-06 | 5.22e-05 |
| POU3F2 | Braak | 0.252 | 5.58e-04 | 1.70e-03 | POU3F2 | CDR | 0.155 | 0.029 | 4.46e-02 |
| POU3F2 | CERAD | 0.313 | 1.17e-04 | 4.61e-04 | ABLIM1 | Braak | 0.339 | 2.35e-06 | 3.09e-05 |
| ABLIM1 | CDR | 0.388 | 1.60e-08 | 4.68e-07 | ABLIM1 | CERAD | 0.431 | 5.21e-08 | 3.36e-06 |
| SMAD4 | Braak | 0.293 | 5.26e-05 | 2.60e-04 | SMAD4 | CDR | 0.295 | 2.36e-05 | 1.05e-04 |
| SMAD4 | CERAD | 0.383 | 1.74e-06 | 2.22e-05 | EP300 | Braak | 0.308 | 2.10e-05 | 1.34e-04 |
| EP300 | CDR | 0.226 | 0.001 | 2.87e-03 | EP300 | CERAD | 0.311 | 1.31e-04 | 4.98e-04 |
| MAP4K5 | Braak | 0.312 | 1.59e-05 | 1.10e-04 | MAP4K5 | CDR | 0.210 | 0.003 | 5.83e-03 |
| MAP4K5 | CERAD | 0.362 | 7.04e-06 | 5.63e-05 | EYA2 | Braak | 0.353 | 8.87e-07 | 1.43e-05 |
| EYA2 | CDR | 0.340 | 9.41e-07 | 8.20e-06 | EYA2 | CERAD | 0.363 | 6.43e-06 | 5.30e-05 |
| MED13 | Braak | 0.171 | 0.020 | 3.37e-02 | MED13 | CDR | 0.160 | 0.024 | 3.90e-02 |
| MED13 | CERAD | 0.302 | 2.06e-04 | 7.10e-04 | SOCS4 | Braak | 0.194 | 0.008 | 1.59e-02 |
| SOCS4 | CDR | 0.205 | 0.004 | 7.01e-03 | SOCS4 | CERAD | 0.326 | 5.83e-05 | 2.76e-04 |
| KANK1 | Braak | 0.300 | 3.31e-05 | 1.90e-04 | KANK1 | CDR | 0.379 | 3.41e-08 | 6.87e-07 |
| KANK1 | CERAD | 0.352 | 1.26e-05 | 8.64e-05 | PDPR | Braak | 0.175 | 0.017 | 3.04e-02 |
| PDPR | CDR | 0.303 | 1.42e-05 | 6.84e-05 | PDPR | CERAD | 0.288 | 4.07e-04 | 1.26e-03 |
| TMEM150A | Braak | 0.325 | 6.42e-06 | 5.87e-05 | TMEM150A | CDR | 0.304 | 1.25e-05 | 6.25e-05 |
| TMEM150A | CERAD | 0.334 | 3.60e-05 | 1.93e-04 | PAPLN | Braak | 0.329 | 4.83e-06 | 4.94e-05 |
| PAPLN | CDR | 0.325 | 2.84e-06 | 1.91e-05 | PAPLN | CERAD | 0.312 | 1.23e-04 | 4.75e-04 |
| EIF2S1 | Braak | -0.297 | 4.24e-05 | 2.18e-04 | EIF2S1 | CDR | -0.319 | 4.59e-06 | 2.75e-05 |
| EIF2S1 | CERAD | -0.272 | 8.85e-04 | 2.33e-03 | SLC25A18 | Braak | 0.385 | 6.64e-08 | 3.39e-06 |
| SLC25A18 | CDR | 0.201 | 0.004 | 8.37e-03 | SLC25A18 | CERAD | 0.282 | 5.59e-04 | 1.62e-03 |
| EFNB1 | Braak | 0.202 | 0.006 | 1.20e-02 | EFNB1 | CDR | 0.362 | 1.54e-07 | 2.06e-06 |
| EFNB1 | CERAD | 0.353 | 1.20e-05 | 8.34e-05 | ZNF395 | Braak | 0.304 | 2.57e-05 | 1.55e-04 |
| ZNF395 | CDR | 0.419 | 7.40e-10 | 6.82e-08 | ZNF395 | CERAD | 0.370 | 4.04e-06 | 4.15e-05 |
| BBX | Braak | 0.210 | 0.004 | 9.37e-03 | BBX | CDR | 0.235 | 8.30e-04 | 1.89e-03 |
| BBX | CERAD | 0.326 | 5.83e-05 | 2.76e-04 | NEK9 | Braak | 0.373 | 1.75e-07 | 4.51e-06 |
| NEK9 | CDR | 0.422 | 5.54e-10 | 5.99e-08 | NEK9 | CERAD | 0.410 | 2.63e-07 | 6.29e-06 |
| MCL1 | Braak | 0.308 | 2.08e-05 | 1.34e-04 | MCL1 | CDR | 0.310 | 8.39e-06 | 4.40e-05 |
| MCL1 | CERAD | 0.308 | 1.52e-04 | 5.57e-04 | FAM107A | Braak | 0.377 | 1.28e-07 | 3.73e-06 |
| FAM107A | CDR | 0.375 | 5.16e-08 | 8.75e-07 | FAM107A | CERAD | 0.429 | 6.37e-08 | 3.42e-06 |
| SOAT1 | Braak | 0.293 | 5.38e-05 | 2.61e-04 | SOAT1 | CDR | 0.272 | 1.05e-04 | 3.51e-04 |
| SOAT1 | CERAD | 0.363 | 6.50e-06 | 5.30e-05 | FMNL3 | Braak | 0.328 | 5.15e-06 | 5.11e-05 |
| FMNL3 | CDR | 0.380 | 3.35e-08 | 6.87e-07 | FMNL3 | CERAD | 0.359 | 8.40e-06 | 6.25e-05 |
| MCMBP | Braak | 0.217 | 0.003 | 7.05e-03 | MCMBP | CDR | 0.271 | 1.09e-04 | 3.61e-04 |
| MCMBP | CERAD | 0.347 | 1.76e-05 | 1.12e-04 | STAG1 | Braak | 0.177 | 0.016 | 2.80e-02 |
| STAG1 | CDR | 0.200 | 0.005 | 8.58e-03 | STAG1 | CERAD | 0.300 | 2.24e-04 | 7.67e-04 |
| CYP2J2 | Braak | 0.268 | 2.30e-04 | 8.23e-04 | CYP2J2 | CDR | 0.301 | 1.61e-05 | 7.58e-05 |
| CYP2J2 | CERAD | 0.280 | 5.92e-04 | 1.68e-03 | BTN3A2 | Braak | 0.246 | 7.32e-04 | 2.11e-03 |
| BTN3A2 | CDR | 0.284 | 4.77e-05 | 1.87e-04 | BTN3A2 | CERAD | 0.333 | 3.82e-05 | 1.98e-04 |
| EFCAB14 | Braak | 0.276 | 1.48e-04 | 5.77e-04 | EFCAB14 | CDR | 0.270 | 1.17e-04 | 3.78e-04 |
| EFCAB14 | CERAD | 0.346 | 1.83e-05 | 1.15e-04 | EIF4EBP2 | Braak | 0.293 | 5.33e-05 | 2.60e-04 |
| EIF4EBP2 | CDR | 0.260 | 2.14e-04 | 5.97e-04 | EIF4EBP2 | CERAD | 0.318 | 8.84e-05 | 3.75e-04 |
| PXN | Braak | 0.351 | 9.67e-07 | 1.48e-05 | PXN | CDR | 0.371 | 6.82e-08 | 1.09e-06 |
| PXN | CERAD | 0.364 | 5.94e-06 | 5.22e-05 | TNIP2 | Braak | 0.251 | 5.60e-04 | 1.70e-03 |
| TNIP2 | CDR | 0.312 | 7.35e-06 | 3.98e-05 | TNIP2 | CERAD | 0.238 | 0.004 | 7.59e-03 |
| TJP1 | Braak | 0.288 | 7.22e-05 | 3.34e-04 | TJP1 | CDR | 0.313 | 7.13e-06 | 3.96e-05 |
| TJP1 | CERAD | 0.360 | 7.86e-06 | 6.11e-05 | KCTD11 | Braak | 0.277 | 1.39e-04 | 5.60e-04 |
| KCTD11 | CDR | 0.334 | 1.45e-06 | 1.18e-05 | KCTD11 | CERAD | 0.335 | 3.55e-05 | 1.92e-04 |
| TTC38 | Braak | 0.232 | 0.001 | 3.75e-03 | TTC38 | CDR | 0.383 | 2.50e-08 | 6.20e-07 |
| TTC38 | CERAD | 0.280 | 6.12e-04 | 1.72e-03 | CRTAP | Braak | 0.414 | 4.99e-09 | 1.07e-06 |

| Gene | Trait | r | p | FDR | Gene | Trait | r | p | FDR |
|---|---|---|---|---|---|---|---|---|---|
| CRTAP | CDR | 0.330 | 1.91e-06 | 1.44e-05 | CRTAP | CERAD | 0.414 | 2.04e-07 | 5.74e-06 |
| YBX1 | Braak | 0.264 | 2.88e-04 | 9.83e-04 | YBX1 | CDR | 0.357 | 2.34e-07 | 2.84e-06 |
| YBX1 | CERAD | 0.337 | 3.06e-05 | 1.73e-04 | TCEA3 | Braak | 0.335 | 3.31e-06 | 4.02e-05 |
| TCEA3 | CDR | 0.324 | 3.04e-06 | 2.02e-05 | TCEA3 | CERAD | 0.330 | 4.55e-05 | 2.27e-04 |
| PBX3 | Braak | 0.314 | 1.38e-05 | 1.00e-04 | PBX3 | CDR | 0.336 | 1.28e-06 | 1.05e-05 |
| PBX3 | CERAD | 0.374 | 3.23e-06 | 3.47e-05 | OLFM1 | Braak | -0.322 | 8.07e-06 | 6.48e-05 |
| OLFM1 | CDR | -0.381 | 2.86e-08 | 6.27e-07 | OLFM1 | CERAD | -0.355 | 1.10e-05 | 7.77e-05 |
| RRAD | Braak | 0.261 | 3.41e-04 | 1.11e-03 | RRAD | CDR | 0.250 | 3.63e-04 | 9.20e-04 |
| RRAD | CERAD | 0.365 | 5.72e-06 | 5.19e-05 | AEN | Braak | 0.303 | 2.85e-05 | 1.68e-04 |
| AEN | CDR | 0.260 | 2.07e-04 | 5.86e-04 | AEN | CERAD | 0.305 | 1.79e-04 | 6.32e-04 |
| PRKCA | Braak | 0.203 | 0.006 | 1.16e-02 | PRKCA | CDR | 0.176 | 0.013 | 2.14e-02 |
| PRKCA | CERAD | 0.318 | 9.00e-05 | 3.79e-04 | FMNL2 | Braak | 0.310 | 1.80e-05 | 1.22e-04 |
| FMNL2 | CDR | 0.232 | 9.63e-04 | 2.16e-03 | FMNL2 | CERAD | 0.313 | 1.15e-04 | 4.61e-04 |
| MGAT1 | Braak | 0.292 | 5.48e-05 | 2.61e-04 | MGAT1 | CDR | 0.325 | 2.86e-06 | 1.91e-05 |
| MGAT1 | CERAD | 0.278 | 6.77e-04 | 1.89e-03 | BHLHE41 | Braak | 0.259 | 3.66e-04 | 1.17e-03 |
| BHLHE41 | CDR | 0.209 | 0.003 | 6.06e-03 | BHLHE41 | CERAD | 0.322 | 7.26e-05 | 3.23e-04 |
| TGIF1 | Braak | 0.354 | 7.87e-07 | 1.34e-05 | TGIF1 | CDR | 0.381 | 2.92e-08 | 6.27e-07 |
| TGIF1 | CERAD | 0.403 | 4.27e-07 | 8.87e-06 | SLFN11 | Braak | 0.291 | 5.84e-05 | 2.77e-04 |
| SLFN11 | CDR | 0.270 | 1.19e-04 | 3.82e-04 | SLFN11 | CERAD | 0.362 | 7.08e-06 | 5.63e-05 |
| ZXDC | Braak | 0.322 | 8.15e-06 | 6.48e-05 | ZXDC | CDR | 0.354 | 3.01e-07 | 3.60e-06 |
| ZXDC | CERAD | 0.337 | 3.10e-05 | 1.73e-04 | ATM | Braak | 0.222 | 0.002 | 5.69e-03 |
| ATM | CDR | 0.194 | 0.006 | 1.11e-02 | ATM | CERAD | 0.303 | 1.94e-04 | 6.75e-04 |
| RXRA | Braak | 0.334 | 3.48e-06 | 4.15e-05 | RXRA | CDR | 0.296 | 2.21e-05 | 1.00e-04 |
| RXRA | CERAD | 0.263 | 0.001 | 3.24e-03 | PLIN5 | Braak | 0.308 | 2.11e-05 | 1.34e-04 |
| PLIN5 | CDR | 0.329 | 2.20e-06 | 1.58e-05 | PLIN5 | CERAD | 0.163 | 0.049 | 6.73e-02 |
| PHKG1 | Braak | 0.281 | 1.08e-04 | 4.66e-04 | PHKG1 | CDR | 0.357 | 2.25e-07 | 2.79e-06 |
| PHKG1 | CERAD | 0.342 | 2.28e-05 | 1.37e-04 | KDSR | Braak | 0.244 | 8.23e-04 | 2.29e-03 |
| KDSR | CDR | 0.178 | 0.012 | 2.02e-02 | KDSR | CERAD | 0.303 | 1.95e-04 | 6.77e-04 |
| NPL | Braak | 0.333 | 3.67e-06 | 4.15e-05 | NPL | CDR | 0.414 | 1.32e-09 | 9.47e-08 |
| NPL | CERAD | 0.407 | 3.32e-07 | 7.64e-06 | DOCK1 | Braak | 0.397 | 2.29e-08 | 1.84e-06 |
| DOCK1 | CDR | 0.319 | 4.40e-06 | 2.73e-05 | DOCK1 | CERAD | 0.352 | 1.30e-05 | 8.75e-05 |
| RAD52 | Braak | 0.273 | 1.69e-04 | 6.46e-04 | RAD52 | CDR | 0.324 | 3.07e-06 | 2.02e-05 |
| RAD52 | CERAD | 0.229 | 0.005 | 9.91e-03 | NEK6 | Braak | 0.297 | 4.20e-05 | 2.18e-04 |
| NEK6 | CDR | 0.340 | 9.12e-07 | 8.20e-06 | NEK6 | CERAD | 0.380 | 2.23e-06 | 2.62e-05 |
| NPC2 | Braak | 0.206 | 0.005 | 1.05e-02 | NPC2 | CDR | 0.287 | 3.97e-05 | 1.60e-04 |
| NPC2 | CERAD | 0.322 | 7.32e-05 | 3.23e-04 | VAMP5 | Braak | 0.197 | 0.007 | 1.44e-02 |
| VAMP5 | CDR | 0.280 | 6.39e-05 | 2.27e-04 | VAMP5 | CERAD | 0.321 | 7.51e-05 | 3.29e-04 |
| FAM167A | Braak | 0.272 | 1.86e-04 | 7.04e-04 | FAM167A | CDR | 0.330 | 1.97e-06 | 1.47e-05 |
| FAM167A | CERAD | 0.392 | 9.64e-07 | 1.46e-05 | HDAC1 | Braak | 0.339 | 2.35e-06 | 3.09e-05 |
| HDAC1 | CDR | 0.440 | 8.13e-11 | 3.87e-08 | HDAC1 | CERAD | 0.325 | 6.08e-05 | 2.83e-04 |
| EVA1B | Braak | 0.258 | 3.93e-04 | 1.24e-03 | EVA1B | CDR | 0.353 | 3.37e-07 | 3.81e-06 |
| EVA1B | CERAD | 0.210 | 0.011 | 1.80e-02 | NUP188 | Braak | 0.286 | 7.92e-05 | 3.55e-04 |
| NUP188 | CDR | 0.313 | 6.76e-06 | 3.82e-05 | NUP188 | CERAD | 0.370 | 4.15e-06 | 4.18e-05 |
| GPNMB | Braak | 0.215 | 0.003 | 7.41e-03 | GPNMB | CDR | 0.130 | 0.067 | 9.40e-02 |
| GPNMB | CERAD | 0.311 | 1.31e-04 | 4.98e-04 | HBP1 | Braak | 0.249 | 6.29e-04 | 1.85e-03 |
| HBP1 | CDR | 0.204 | 0.004 | 7.48e-03 | HBP1 | CERAD | 0.337 | 3.15e-05 | 1.73e-04 |
| AMOT | Braak | 0.356 | 6.67e-07 | 1.22e-05 | AMOT | CDR | 0.326 | 2.63e-06 | 1.80e-05 |
| AMOT | CERAD | 0.413 | 2.13e-07 | 5.74e-06 | MYLK | Braak | 0.270 | 2.01e-04 | 7.52e-04 |
| MYLK | CDR | 0.333 | 1.60e-06 | 1.26e-05 | MYLK | CERAD | 0.369 | 4.46e-06 | 4.35e-05 |
| SOX13 | Braak | 0.415 | 4.55e-09 | 1.07e-06 | SOX13 | CDR | 0.381 | 2.88e-08 | 6.27e-07 |
| SOX13 | CERAD | 0.364 | 6.04e-06 | 5.22e-05 | CDK2AP1 | Braak | 0.314 | 1.42e-05 | 1.00e-04 |
| CDK2AP1 | CDR | 0.284 | 4.80e-05 | 1.87e-04 | CDK2AP1 | CERAD | 0.334 | 3.67e-05 | 1.94e-04 |
| TAB2 | Braak | 0.286 | 8.03e-05 | 3.57e-04 | TAB2 | CDR | 0.293 | 2.69e-05 | 1.16e-04 |
| TAB2 | CERAD | 0.359 | 8.32e-06 | 6.25e-05 | ACSS1 | Braak | 0.313 | 1.44e-05 | 1.00e-04 |
| ACSS1 | CDR | 0.279 | 6.58e-05 | 2.31e-04 | ACSS1 | CERAD | 0.312 | 1.24e-04 | 4.75e-04 |
| SMAD9 | Braak | 0.247 | 7.08e-04 | 2.05e-03 | SMAD9 | CDR | 0.264 | 1.67e-04 | 4.97e-04 |
| SMAD9 | CERAD | 0.395 | 7.68e-07 | 1.27e-05 | ARHGEF10 | Braak | 0.358 | 5.99e-07 | 1.17e-05 |
| ARHGEF10 | CDR | 0.334 | 1.51e-06 | 1.20e-05 | ARHGEF10 | CERAD | 0.392 | 9.81e-07 | 1.46e-05 |
| NAGLU | Braak | 0.296 | 4.38e-05 | 2.21e-04 | NAGLU | CDR | 0.266 | 1.51e-04 | 4.62e-04 |
| NAGLU | CERAD | 0.316 | 1.01e-04 | 4.13e-04 | WASF2 | Braak | 0.311 | 1.70e-05 | 1.16e-04 |
| WASF2 | CDR | 0.354 | 3.11e-07 | 3.65e-06 | WASF2 | CERAD | 0.397 | 6.74e-07 | 1.20e-05 |
| IRF9 | Braak | 0.320 | 9.20e-06 | 7.18e-05 | IRF9 | CDR | 0.374 | 5.69e-08 | 9.40e-07 |
| IRF9 | CERAD | 0.345 | 1.95e-05 | 1.21e-04 | COPZ2 | Braak | 0.319 | 1.01e-05 | 7.77e-05 |
| COPZ2 | CDR | 0.390 | 1.33e-08 | 4.09e-07 | COPZ2 | CERAD | 0.438 | 3.17e-08 | 2.70e-06 |
| SOGA1 | Braak | 0.276 | 1.47e-04 | 5.77e-04 | SOGA1 | CDR | 0.313 | 6.67e-06 | 3.80e-05 |
| SOGA1 | CERAD | 0.293 | 3.19e-04 | 1.04e-03 | BAZ2B | Braak | 0.242 | 9.29e-04 | 2.53e-03 |
| BAZ2B | CDR | 0.247 | 4.40e-04 | 1.08e-03 | BAZ2B | CERAD | 0.358 | 9.11e-06 | 6.67e-05 |
| CPT1A | Braak | 0.313 | 1.43e-05 | 1.00e-04 | CPT1A | CDR | 0.262 | 1.82e-04 | 5.33e-04 |
| CPT1A | CERAD | 0.246 | 0.003 | 5.73e-03 | TNS3 | Braak | 0.356 | 6.67e-07 | 1.22e-05 |
| TNS3 | CDR | 0.376 | 4.67e-08 | 8.37e-07 | TNS3 | CERAD | 0.329 | 4.93e-05 | 2.44e-04 |
| SERPING1 | Braak | 0.235 | 0.001 | 3.29e-03 | SERPING1 | CDR | 0.280 | 6.24e-05 | 2.23e-04 |
| SERPING1 | CERAD | 0.307 | 1.58e-04 | 5.73e-04 | HLA-DPB1 | Braak | 0.236 | 0.001 | 3.24e-03 |
| HLA-DPB1 | CDR | 0.277 | 7.72e-05 | 2.67e-04 | HLA-DPB1 | CERAD | 0.370 | 4.26e-06 | 4.23e-05 |
| HSPB1 | Braak | 0.280 | 1.12e-04 | 4.73e-04 | HSPB1 | CDR | 0.316 | 5.62e-06 | 3.23e-05 |
| HSPB1 | CERAD | 0.292 | 3.38e-04 | 1.09e-03 | GYPC | Braak | 0.327 | 5.82e-06 | 5.60e-05 |
| GYPC | CDR | 0.431 | 2.29e-10 | 3.87e-08 | GYPC | CERAD | 0.448 | 1.42e-08 | 1.83e-06 |
| PRDX6 | Braak | 0.231 | 0.002 | 3.95e-03 | PRDX6 | CDR | 0.266 | 1.52e-04 | 4.63e-04 |
| PRDX6 | CERAD | 0.323 | 6.91e-05 | 3.14e-04 | LCAT | Braak | 0.363 | 3.97e-07 | 8.53e-06 |

| Gene | Trait | r | p | FDR | Gene | Trait | r | p | FDR |
|---|---|---|---|---|---|---|---|---|---|
| LCAT | CDR | 0.362 | 1.57e-07 | 2.06e-06 | LCAT | CERAD | 0.367 | 5.08e-06 | 4.70e-05 |
| ATL3 | Braak | 0.244 | 8.23e-04 | 2.29e-03 | ATL3 | CDR | 0.271 | 1.12e-04 | 3.67e-04 |
| ATL3 | CERAD | 0.328 | 5.25e-05 | 2.56e-04 | POLDIP3 | Braak | 0.298 | 3.87e-05 | 2.08e-04 |
| POLDIP3 | CDR | 0.319 | 4.43e-06 | 2.73e-05 | POLDIP3 | CERAD | 0.364 | 6.15e-06 | 5.22e-05 |
| TP53INP1 | Braak | 0.343 | 1.77e-06 | 2.48e-05 | TP53INP1 | CDR | 0.341 | 8.73e-07 | 8.04e-06 |
| TP53INP1 | CERAD | 0.420 | 1.31e-07 | 5.63e-06 | SH3BP4 | Braak | 0.300 | 3.35e-05 | 1.91e-04 |
| SH3BP4 | CDR | 0.312 | 7.20e-06 | 3.96e-05 | SH3BP4 | CERAD | 0.332 | 4.02e-05 | 2.04e-04 |
| ATP1B2 | Braak | 0.299 | 3.60e-05 | 1.99e-04 | ATP1B2 | CDR | 0.172 | 0.015 | 2.57e-02 |
| ATP1B2 | CERAD | 0.316 | 9.89e-05 | 4.11e-04 | GPRC5C | Braak | 0.184 | 0.012 | 2.26e-02 |
| GPRC5C | CDR | 0.309 | 9.23e-06 | 4.75e-05 | GPRC5C | CERAD | 0.305 | 1.74e-04 | 6.20e-04 |
| EEF1A2 | Braak | -0.264 | 2.91e-04 | 9.83e-04 | EEF1A2 | CDR | -0.291 | 3.12e-05 | 1.32e-04 |
| EEF1A2 | CERAD | -0.312 | 1.21e-04 | 4.69e-04 | HLA-E | Braak | 0.222 | 0.002 | 5.69e-03 |
| HLA-E | CDR | 0.309 | 9.39e-06 | 4.76e-05 | HLA-E | CERAD | 0.309 | 1.45e-04 | 5.40e-04 |
| KSR1 | Braak | 0.261 | 3.38e-04 | 1.11e-03 | KSR1 | CDR | 0.303 | 1.40e-05 | 6.81e-05 |
| KSR1 | CERAD | 0.264 | 0.001 | 3.08e-03 | PPFIBP2 | Braak | 0.329 | 4.82e-06 | 4.94e-05 |
| PPFIBP2 | CDR | 0.256 | 2.62e-04 | 7.05e-04 | PPFIBP2 | CERAD | 0.344 | 2.07e-05 | 1.26e-04 |
| TRIOBP | Braak | 0.388 | 4.97e-08 | 2.91e-06 | TRIOBP | CDR | 0.319 | 4.42e-06 | 2.73e-05 |
| TRIOBP | CERAD | 0.415 | 1.83e-07 | 5.74e-06 | CABP1 | Braak | -0.264 | 2.93e-04 | 9.83e-04 |
| CABP1 | CDR | -0.329 | 2.10e-06 | 1.53e-05 | CABP1 | CERAD | -0.364 | 6.01e-06 | 5.22e-05 |
| STON2 | Braak | 0.329 | 5.11e-06 | 5.11e-05 | STON2 | CDR | 0.409 | 2.17e-09 | 1.27e-07 |
| STON2 | CERAD | 0.429 | 6.22e-08 | 3.42e-06 | GPT2 | Braak | 0.323 | 7.43e-06 | 6.27e-05 |
| GPT2 | CDR | 0.291 | 3.12e-05 | 1.32e-04 | GPT2 | CERAD | 0.283 | 5.16e-04 | 1.52e-03 |
| SERINC5 | Braak | 0.223 | 0.002 | 5.57e-03 | SERINC5 | CDR | 0.303 | 1.40e-05 | 6.81e-05 |
| SERINC5 | CERAD | 0.313 | 1.17e-04 | 4.61e-04 | PPP1R18 | Braak | 0.261 | 3.40e-04 | 1.11e-03 |
| PPP1R18 | CDR | 0.334 | 1.50e-06 | 1.20e-05 | PPP1R18 | CERAD | 0.261 | 0.001 | 3.49e-03 |
| ARPC1B | Braak | 0.260 | 3.63e-04 | 1.17e-03 | ARPC1B | CDR | 0.312 | 7.18e-06 | 3.96e-05 |
| ARPC1B | CERAD | 0.265 | 0.001 | 3.01e-03 | RHOC | Braak | 0.202 | 0.006 | 1.21e-02 |
| RHOC | CDR | 0.364 | 1.31e-07 | 1.83e-06 | RHOC | CERAD | 0.289 | 3.95e-04 | 1.23e-03 |
| PLCG2 | Braak | 0.286 | 8.15e-05 | 3.57e-04 | PLCG2 | CDR | 0.336 | 1.25e-06 | 1.05e-05 |
| PLCG2 | CERAD | 0.326 | 5.76e-05 | 2.76e-04 | P2RX7 | Braak | 0.324 | 7.12e-06 | 6.26e-05 |
| P2RX7 | CDR | 0.248 | 4.17e-04 | 1.04e-03 | P2RX7 | CERAD | 0.247 | 0.003 | 5.49e-03 |
| XAF1 | Braak | 0.285 | 8.39e-05 | 3.65e-04 | XAF1 | CDR | 0.329 | 2.21e-06 | 1.58e-05 |
| XAF1 | CERAD | 0.319 | 8.60e-05 | 3.71e-04 | DDAH2 | Braak | 0.303 | 2.88e-05 | 1.69e-04 |
| DDAH2 | CDR | 0.352 | 3.62e-07 | 4.02e-06 | DDAH2 | CERAD | 0.361 | 7.46e-06 | 5.86e-05 |
| CAPG | Braak | 0.176 | 0.016 | 2.88e-02 | CAPG | CDR | 0.353 | 3.22e-07 | 3.71e-06 |
| CAPG | CERAD | 0.340 | 2.66e-05 | 1.54e-04 | | | | | |

### References


[1] Soneson, C. (2014). compcodeR—an R package for benchmarking differential expression methods for RNA-seq data. Bioinformatics, 30(17), 2517–2518.

[2] Allen, M. et al. (2016). Human whole genome genotype and transcriptome data for Alzheimer's and other neurodegenerative diseases. Scientific Data, 3, 160089.

[3] Wang, M. et al. (2018). The Mount Sinai cohort of large-scale genomic, transcriptomic and proteomic data in Alzheimer's disease. Scientific Data, 5, 180185.

[4] Patro, R. et al. (2017). Salmon provides fast and bias-aware quantification of transcript expression. Nature Methods, 14, 417–419.

[5] Love, M.I., Huber, W. and Anders, S. (2014). Moderated estimation of fold change and dispersion for RNA-seq data with DESeq2. Genome Biology, 15, 550.

[6] Robinson, M.D., McCarthy, D.J. and Smyth, G.K. (2010). edgeR: a Bioconductor package for differential expression analysis of digital gene expression data. Bioinformatics, 26(1), 139–140.

[7] Li, H.D. et al. (2021). Integrative functional genomic analysis of intron retention in human and mouse brain with Alzheimer's disease. Alzheimer's & Dementia, 17(6), 984–1004.

[8] Cerezo, M. et al. (2025). The NHGRI-EBI GWAS Catalog: standards for reusability, sustainability and diversity. Nucleic Acids Research, 53(D1), D998–D1005.

[9] Hagberg, A.A., Schult, D.A. and Swart, P.J. (2008). Exploring network structure, dynamics, and function using NetworkX. Proceedings of the 7th Python in Science Conference, 11–15.

[10] Lin, C.X. et al. (2022). An integrated brain-specific network identifies genes associated with neuropathologic and clinical traits of Alzheimer's disease. Briefings in Bioinformatics, 23(1), bbab522.
